# Strains used in whole organism *Plasmodium falciparum* vaccine trials differ in genome structure, sequence, and immunogenic potential

**DOI:** 10.1101/684175

**Authors:** Kara A. Moser, Elliott F. Drábek, Ankit Dwivedi, Jonathan Crabtree, Emily M. Stucke, Antoine Dara, Zalak Shah, Matthew Adams, Tao Li, Priscila T. Rodrigues, Sergey Koren, Adam M. Phillippy, Amed Ouattara, Kirsten E. Lyke, Lisa Sadzewicz, Luke J. Tallon, Michele D. Spring, Krisada Jongsakul, Chanthap Lon, David L. Saunders, Marcelo U. Ferreira, Myaing M. Nyunt, Miriam K. Laufer, Mark A. Travassos, Robert W. Sauerwein, Shannon Takala-Harrison, Claire M. Fraser, B. Kim Lee Sim, Stephen L. Hoffman, Christopher V. Plowe, Joana C. Silva

## Abstract

**Background:** *Plasmodium falciparum* (Pf) whole-organism sporozoite vaccines have provided excellent protection against controlled human malaria infection (CHMI) and naturally transmitted heterogeneous Pf in the field. Initial CHMI studies showed significantly higher durable protection against homologous than heterologous strains, suggesting the presence of strain-specific vaccine-induced protection. However, interpretation of these results and understanding of their relevance to vaccine efficacy (VE) have been hampered by the lack of knowledge on genetic differences between vaccine and CHMI strains, and how these strains are related to parasites in malaria endemic regions.

**Methods:** Whole genome sequencing using long-read (Pacific Biosciences) and short-read (Illumina) sequencing platforms was conducted to generate *de novo* genome assemblies for the vaccine strain, NF54, and for strains used in heterologous CHMI (7G8 from Brazil, NF166.C8 from Guinea, and NF135.C10 from Cambodia). The assemblies were used to characterize sequence polymorphisms and structural variants in each strain relative to the reference Pf 3D7 (a clone of NF54) genome. Strains were compared to each other and to clinical isolates from South America, Sub-Saharan Africa, and Southeast Asia.

**Results:** While few variants were detected between 3D7 and NF54, we identified tens of thousands of variants between NF54 and the three heterologous strains both genome-wide and within regulatory and immunologically important regions, including in pre-erythrocytic antigens that may be key for sporozoite vaccine-induced protection. Additionally, these variants directly contribute to diversity in immunologically important regions of the genomes as detected through *in silico* CD8^+^ T cell epitope predictions. Of all heterologous strains, NF135.C10 consistently had the highest number of unique predicted epitope sequences when compared to NF54, while NF166.C8 had the lowest. Comparison to global clinical isolates revealed that these four strains are representative of their geographic region of origin despite long-term culture adaptation; of note, NF135.C10 is from an admixed population, and not part of recently-formed drug resistant subpopulations present in the Greater Mekong Sub-region.

**Conclusions:** These results are assisting the interpretation of VE of whole-organism vaccines against homologous and heterologous CHMI, and may be useful in informing the choice of strains for inclusion in region-specific or multi-strain vaccines.

## Introduction

The flattening levels of mortality and morbidity due to malaria in recent years (1), which follow a decade in which malaria mortality was cut in half (2), highlight the pressing need for new tools to control the disease. A highly efficacious vaccine against *Plasmodium falciparum*, the deadliest malaria parasite, would be a critical development for control and elimination efforts. Several variations of a highly promising pre-erythrocytic, whole organism malaria vaccine based on *P. falciparum* sporozoites (PfSPZ) are under development, all based on the same *P. falciparum* strain, NF54 (3), thought to be of West African origin, and which use different mechanisms for attenuation of PfSPZ. Of these vaccine candidates, Sanaria® PfSPZ Vaccine, based on radiation-attenuated sporozoites, has progressed furthest in clinical trial testing (4–10). Other whole-organism vaccine candidates, including chemo-attenuated (Sanaria® PfSPZ-CVac), transgenic and genetically attenuated sporozoites, are in earlier stages of development (11–13).

PfSPZ Vaccine showed 100% short-term protection against homologous controlled human malaria infection (CHMI) in an initial phase 1 clinical trial (6), and subsequent trials have confirmed that high levels of protection can be achieved against both short-term (8) and long-term (7) homologous CHMI. However, depending on the immunization regimen, sterile protection can be significantly lower (8-83%) against heterologous CHMI using the 7G8 Brazilian clone (8, 9), and against infection in malaria-endemic regions with intense seasonal malaria transmission (29% and 52% by proportional and time to event analysis, respectively) (10). Heterologous CHMI in chemoprophylaxis with sporozoite (CPS) studies, in which immunization is by infected mosquito bite of individuals undergoing malaria chemoprophylaxis, have been conducted with NF135.C10 from Cambodia (14) and NF166.C8 from Guinea (15), and have had lower efficacy than against homologous CHMI (16, 17). One explanation for the lower efficacy seen against heterologous *P. falciparum* strains is the extensive genetic diversity in this parasite species, which is particularly high in genes encoding antigens (18), which combined with low vaccine efficacy against non-vaccine alleles (19–21) reduces overall protective efficacy and complicates the design of broadly efficacious vaccines (22, 23). The lack of a detailed genomic characterization of the *P. falciparum* strains used in CHMI studies, and the unknown genetic basis of the parasite targets of PfSPZ Vaccine- and PfSPZ CVac-induced protection, have precluded a conclusive statement regarding the cause(s) of variable vaccine efficacy outcomes.

The current PfSPZ Vaccine strain, NF54, was isolated from a patient in the Netherlands who had never left the country and is considered a case of “airport malaria”; the exact origin of NF54 is unknown (3), but thought to be from Africa (24, 25). NF54 is also the isolate from which the *P. falciparum* 3D7 reference strain was cloned (26), and hence, despite having been separated in culture for over 30 years, NF54 and 3D7 are assumed to be genetically identical, and 3D7 is often used in homologous CHMI (6, 8). Several issues hinder the interpretation of both homologous and heterologous CHMI experiments conducted to date. It remains to be confirmed that 3D7 has remained genetically identical to NF54 genome-wide, or that the two are at least identical immunogenically. Indeed, NF54 and 3D7 have several reported phenotypic differences when grown in culture, including the variable ability to produce gametocytes (27). In addition, 7G8, NF166.C8, and NF135.C10 have not been rigorously compared to each other or to NF54 to confirm that they are adequate heterologous strains, even though they do appear to have distinct infectivity phenotypes when used as CHMI strains (15). While the entire sporozoite likely offers multiple immunological targets, no high-confidence correlates of protection currently exist. In part because of the difficulty of studying hepatic parasite forms and their gene expression profiles in humans, it remains unclear which parasite proteins are recognized by the human immune system during that stage, and elicit protection, upon immunization with PfSPZ vaccines. Both humoral and cell-mediated responses have been associated with protection against homologous CHMI (6, 7), although studies in rodents and non-human primates point to a requirement for cell-mediated immunity (specifically through tissue-resident CD8^+^ T cells) in long-term protection (5,9,28,29). *In silico* identification of CD8^+^ epitopes in all strains could highlight critical differences of immunological significance between strains. Finally, heterologous CHMI results cannot be a reliable indicator of efficacy against infection in field settings unless the CHMI strains used are characteristic of the geographic region from which they originate. These issues could impact the use of homologous and heterologous CHMI, and the choice of strains for these studies, to predict the efficacy of PfSPZ-based vaccines in the field (30).

These knowledge gaps can be addressed through a rigorous description and comparison of the genome sequence of these strains. High-quality *de novo* assemblies allow characterization of genome composition and structure, as well as the identification of genetic differences between strains. However, the high AT content and repetitive nature of the *P. falciparum* genome greatly complicates genome assembly methods (31). Recently, long-read sequencing technologies have been used to overcome some of these assembly challenges, as was shown with assemblies for 3D7, 7G8, and several other culture-adapted *P. falciparum* strains generated using Pacific Biosciences (PacBio) technology (32–34). However, NF166.C8 and NF135.C10 still lack whole-genome assemblies; in addition, while an assembly for 7G8 is available (33), it is important to characterize the specific 7G8 clone used in heterozygous CHMI, from Sanaria’s working bank, as strains can undergo genetic changes over time in culture (35). Here, reference assemblies for NF54, 7G8, NF166.C8, and NF135.C10 (hereafter referred to as PfSPZ strains) were generated using approaches to take advantage of the resolution power of long-read sequencing data and the low error rate of short-read sequencing platforms. These *de novo* assemblies allowed for the thorough genetic and genomic characterization of the PfSPZ strains and will aid in the interpretation of results from CHMI studies.

## Materials and Methods

### Study design and samples

This study characterized and compared the genomes of four *P. falciparum* strains used in whole organism malaria vaccines and controlled human malaria infections using a combination of long- and short-read whole genome sequencing platforms (see below). In addition, these strains were compared to *P. falciparum* clinical isolates collected from patients in malaria-endemic regions globally, using short-read whole genome sequencing data. Genetic material for the four PfSPZ strains were provided by Sanaria, Inc. Clinical *P. falciparum* isolates from Brazil, Mali, Malawi, Myanmar, Thailand, Laos, and Cambodia were collected between 2009 and 2016 from cross-sectional surveys of malaria burden, longitudinal studies of malaria incidence, and drug efficacy studies done in collaboration with the Division of Malaria Research at the University of Maryland Baltimore, or were otherwise provided by collaborators (**Supplemental Dataset 1**). All samples met the inclusion criteria of the initial study protocol with prior approval from the local ethical review board. Parasite genomic sequencing and analyses were undertaken after prior approval of the University of Maryland School of Medicine Institutional Review Board. These isolates were obtained by venous blood draws; almost all samples were processed using leukocyte depletion methods to improve the parasite-to-human ratio before sequencing. The exceptions were samples from Brazil and Malawi, which were not leukocyte depleted upon collection. These samples underwent a selective whole genome amplification step before sequencing, modified from (36) (the main modification being a DNA dilution and filtration step using vacuum filtration prior to selective whole genome amplification). Cambodia isolates had been sequenced for a previous study (37). In addition to these samples, samples for which whole genome short-read sequencing has previously been generated were obtained from NCBI’s Short Read Archive (SRA) to supplement malaria-endemic regions not represented in our data set (Peru, Columbia, French Guiana, Guinea, Papua New Guinea) and regions where PfSPZ trials are ongoing (Burkina Faso, Kenya, and Tanzania) (**Supplemental Dataset 1**).

### Whole genome sequencing

Genetic material for whole genome sequencing of the PfSPZ strains was generated from a cryovial of each strain’s cell bank with the following identifiers: NF54 Working Cell Bank (WCB): SAN02-073009; 7G8 WCB: SAN02-021214; NF135.C10 WCB: SAN07-010410; NF166.C8 Mother Cell Bank: SAN30-020613. Each cryovial was thawed and maintained in human O+ red blood cells (RBCs) at 2% hematocrit (Hct) in complete growth medium (RPMI 1649 with L-glutamine and 25 mM HEPES supplemented with 10% human O+ serum and hypoxanthine) in a 6-well plate in 5% O2, 5% CO2 and 90% N2 at 37°C. The cultures were then further expanded by adding fresh RBCs every 3-4 days and increased culture hematocrit (Hct) to 5% Hct using a standard method (38). The complete growth medium was replaced daily. When the PfSPZ strain culture volume reached 300-400 mL and a parasitemia of more than 1.5%, the culture suspensions were collected and the parasitized RBCs were pelleted down by centrifugation at 1800 rpm for 5 min. Aliquots of 0.5 mL per cryovial of the parasitized RBCs were stored at −80°C prior to extraction of genomic DNA. Genomic DNA was extracted using the Qiagan Blood DNA Midi Kit (Valencia, CA, USA). Pacific Biosciences (PacBio) sequencing was done for each PfSPZ strain. Total DNA was prepared for PacBio sequencing using the DNA Template Prep Kit 2.0 (Pacific Biosciences, Menlo Park, CA). DNA was fragmented with the Covaris E210, and the fragments were size selected to include those >15 Kbp in length. Libraries were prepared per the manufacturer’s protocol. Four SMRT cells were sequenced per library, using P6C4 chemistry and a 120-minute movie on the PacBio RS II (Pacific Biosystems, Menlo Park, CA).

Short-read sequencing was done for each PfSPZ strain and for our collection of clinical isolates using the Illumina HiSeq 2500 or 4000 platforms. Prepared genomic DNA, extracted from cultured parasites, leukocyte-depleted samples, or from samples that underwent sWGA (see above), was used to construct DNA libraries for sequencing on the Illumina platform using the KAPA Library Preparation Kit (Kapa Biosystems, Woburn, MA). DNA was fragmented with the Covaris E210 or E220 to ∼200 bp. Libraries were prepared using a modified version of manufacturer’s protocol. The DNA was purified between enzymatic reactions and the size selection of the library was performed with AMPure XT beads (Beckman Coulter Genomics, Danvers, MA). When necessary, a PCR amplification step was performed with primers containing an index sequence of six nucleotides in length. Libraries were assessed for concentration and fragment size using the DNA High Sensitivity Assay on the LabChip GX (Perkin Elmer, Waltham, MA). Library concentrations were also assessed by qPCR using the KAPA Library Quantification Kit (Complete, Universal) (Kapa Biosystems, Woburn, MA). The libraries were pooled and sequenced on a 100 bp paired-end Illumina HiSeq 2500 or 4000 run (Illumina, San Diego, CA).

### Assembly generation and characterization of PfSPZ strains

Canu (v1.3) (39) was used to correct and assemble the PacBio reads (corMaxEvidenceErate=0.15 for AT-rich genomes, and using default parameters otherwise). To optimize downstream assembly correction processes and parameters, the percentage of total differences (both in bp and by proportion of the 3D7 genome not captured by the NF54 assembly) between the NF54 assembly and the 3D7 reference (PlasmoDBv24) was calculated after each round of correction. Quiver (smrtanalysis v2.3) (40) was run iteratively with default parameters to reach a (stable) maximum reduction in percent differences between the two genomes and the assemblies were further corrected with Illumina data using Pilon (v1.13) (41) with the following parameters: --fixbases, -- mindepth 5, --K 85, --minmq 0, and --minqual 35.

Assemblies were compared to the 3D7 reference (PlasmoDBv24) using MUMmer’s nucmer (42), and the show-snps function was used to generate a list of SNPs and small (< 50 bp) indels between assemblies. Coding and non-coding variants were classified by comparing the show-snps output with the 3D7 gff3 file using custom scripts. Structural variants, defined as indels, deletions, and tandem or repeat expansion and contractions each greater than 50 bp in length were identified using the nucmer-based Assemblytics tool (43) (unique anchor length: 1 Kbp). Translocations were identified by eye through inspection of mummerplots. Translocations and other large structural variants were confirmed using read-mapping data of the PacBio reads against either the 3D7 reference genome or the respective PfSPZ strain assembly using the NGLMR aligner (44) and visualized using Ribbon (45). Reconstructed exon 1 sequences for *var* genes, encoding *P. falciparum* erythrocyte membrane protein 1 (PfEMP1) antigens, for each PfSPZ strain were recovered using the ETHA package (46). As a check for *var* exon 1 sequences that were missed during the generation of the strain’s assembly, a targeted read capture and assembly approach was done using a strain’s Illumina data, wherein *var*-like reads for each PfSPZ strains were identified by mapping reads against a database of known *var* exon 1 sequences (47) using bowtie2 (48). Reads that mapped to a known exon 1 sequence were then assembled with Spades (v3.9.0) (49), and the assembled products were blasted against the PacBio reads to identify if they were truly missed exon 1 sequences by the *de novo* assembly process, or if instead they were chimeras reconstructed by the targeted assembly process. To identify homologs of the 61 *var* exon 1 sequences found in the 3D7 genome, each PfSPZ strain’s set of exon 1 amino acid sequences were compared to the 3D7 set using blastp.

### In silico MHC I epitope predictions

Given the reported importance of CD8^+^ T cell responses towards immunity to whole sporozoites, MHC class 1 epitopes of length 9 amino acids (9mers) were predicted with NetMHCpan (v3.0) (50) for each PfSPZ strain, using inferred amino acid sequences in each genome. HLA types common to African countries where PfSPZ or PfSPZ-CVac trials are ongoing were used for epitope predictions based on frequencies in the Allele Frequency Net Database (51) or from the literature (52, 53) (**Table S1**). Shared epitopes between NF54 and the three heterologous PfSPZ strains were calculated by first identifying epitopes in each gene, and then removing duplicate epitope sequence entries (caused by recognition by multiple HLA types). Identical epitope sequences that were identified in two or more genes were treated as distinct epitope entries, and all unique epitope-gene combinations were included when calculating the number of shared epitopes between strains.

Epitopes were predicted in several gene sets. The first was all protein-coding genes in which no premature stop codons were detected across all four PfSPZ strains (n=4,814). The second was a subset of the above that are expressed as proteins during the sporozoite stage of the parasite in the mosquito salivary glands, as supported by proteomic data (n=1,974) (54, 55). Lastly, a list of pre-erythrocytic genes of interest was generated, either from a review of the literature, or genes whose products were recognized by sera from protected vaccinees of whole organism malaria vaccine trials (both PfSPZ and PfSPZ-CVac) (n=42) (11, 56). These gene sequences were inspected manually in each assembly, and corrected for any remaining PacBio sequencing error, particularly one bp indels in homopolymer runs. Even though the 42 genes listed above were detected through antibody responses, many have also been shown to have T cell epitopes, such as CSP and LSA-1.

### Read mapping and SNP calling

For the full collection of clinical isolates that had whole genome short-read sequencing data (generated either at IGS or downloaded from SRA), reads were aligned to the 3D7 reference genome (PlasmoDBv24) using bowtie2 (v2.2.4) (48). Samples with less than 10 million reads mapping to the reference were excluded, as samples with less than this amount had reduced coverage across the genome. Bam files were processed according to GATK’s Best Practices documentation (57–59). Joint SNP calling was done using Haplotype Caller (v4.0). Because clinical samples may be polyclonal (that is, more than one parasite strain may be present), diploid calls were initially allowed, followed by calling the major allele at positions with heterozygous calls. If the major allele was supported by >70% of reads at a heterozygous position, the major allele was assigned as the allele at that position (otherwise, the genotype was coded as missing). Additional hard filtering was done to remove potential false positives based on the following filter: DP < 12 || QUAL < 50 || FS > 14.5 || MQ < 20, and variants were further filtered to remove those for which the non-reference allele was not present in at least three samples (frequency less than ∼0.5%), and those with more than 10% missing genotype values across all samples.

### Principal Coordinates Analyses and Admixture Analyses

A matrix of pairwise genetic distances was constructed from biallelic non-synonymous SNPs identified from the above pipeline (n=31,761) across all samples (n=654) using a custom Python script, and principal coordinates analyses (PCoAs) were done to explore population structure using cmdscale in R. Additional population structure analyses were done using Admixture (v1.3) (60) on two separate data sets: South America and Africa clinical isolates plus NF54, NF166.C8, and 7G8 (n=461), and Southeast Asia and Oceania plus NF135.C10 (n=193). The data sets were additionally pruned for sites in linkage disequilibrium (window size of 20 Kbp, window step of 2 Kbp, R^2^ ≥0.1). The final South America/Africa and Southeast Asia/Oceania data set used for the admixture analysis consisted of 16,802 and 5,856 SNPs, respectively. The number of populations, *K*, was tested for values between *K*=1 to *K*=15 and run with 10 replicates for each *K*. For each population, the cross-validation (CV) error from the replicate with the highest log-likelihood value was plotted, and the *K* with the lowest CV value was chosen as the final *K*.

To compare subpopulations identified in our Southeast Asia/Oceania admixture analysis with previously described ancestral, resistant, and admixed subpopulations from Cambodia (61), the above non-synonymous SNP set was used before pruning for LD (n=11,943), and was compared to a non-synonymous SNP dataset (n=21,257) from 167 samples used by Dwivedi *et al.* (62) to describe eight Cambodian subpopulations, in an analysis that included a subset of samples used by Miotto *et al.* (61) (who first characterized the population structure in Cambodia). There were 5,881 shared non-synonymous SNPs between the two datasets, 1,649 of which were observed in NF135.C10. A pairwise genetic distance matrix (estimated as the proportion of base-pair differences between pairs of samples, not including missing genotypes) was generated from the 5,881 shared SNP set, and a dendrogram was built using Ward minimum variance methods in R (Ward.D2 option of the hclust function).

## Results

### Generation of assemblies and validation of the assembly protocol

To characterize genome-wide structural and genetic diversity of the PfSPZ strains, genome assemblies were generated *de novo* using whole genome long-read (PacBio) and short-read (Illumina) sequence data (Methods; **Table S2**; **Table S3**). Taking advantage of the parent isolate-clone relationship between NF54 and 3D7, we used NF54 as a test case to derive the assembly protocol, by adopting, at each step, approaches that minimized the difference to 3D7. The raw assembly of the NF54 genome, while containing 99.9% of the 3D7 genome in 30 contigs, showed ∼78K sequence variants relative to 3D7 (**Figure S1**). Many of these differences were small (1–3) base pair (bp) indels (insertions or deletions), particularly in AT-rich regions and homopolymer runs, strongly indicative of remaining sequencing errors (63). To maximize error removal with existing tools, Quiver was run iteratively on the NF54 assembly; two consecutive Quiver runs consistently and substantially reduced the number of differences between NF54 and 3D7 (**Figure S1**). Illumina reads from NF54 were used to further polish the assembly using Pilon. The resulting NF54 assembly is 23.4 Mb in length (**Figure 1**) and, when compared to the 3D7 reference, contains 8,396 indels and 1,386 single nucleotide polymorphisms (SNPs), a rate of ∼59 SNPs per million base pairs, the vast majority of which occurred in noncoding regions (**Table 1**). In fact, only 17 non-synonymous mutations were detected in 15 intact protein-coding single-copy genes between the two genomes (**Supplemental Dataset 2**). We applied the same assembly protocol to the remaining PfSPZ strains. The resulting assemblies for NF166.C8, 7G8, and NF135.C10 are structurally very complete, with the 14 nuclear chromosomes represented by 30, 20, and 19 nuclear contigs, respectively, with each chromosome in the 3D7 reference represented by one to three contigs (**Figure 1**). Several shorter contigs in NF54 (67,501 bps total), NF166.C8 (224,502 bps total), and NF135.C10 (80,944 bps total) could not be unambiguously assigned to a homologous segment in the 3D7 reference genome; gene annotation showed that these contigs mostly contain members of multi-gene families, and therefore are likely part of sub-telomeric regions. The cumulative lengths of the four assemblies ranged from 22.8 Mbp to 23.5 Mbp (**Table 1**), indicating variation in genome size among *P. falciparum* strains. In particular, the 7G8 assembly was several hundred thousand base-pairs smaller than the other three assemblies. To confirm that this was not an assembly error, we compared 7G8 to a previously published 7G8 PacBio-based assembly (PlasmoDBv41) (33). The two assemblies were extremely close in overall genome structure, differing only by 2 Kbp in cumulative length, and also shared a very similar number of SNP and small indel variants relative to 3D7 (**Table S4**).

**Figure 1:**
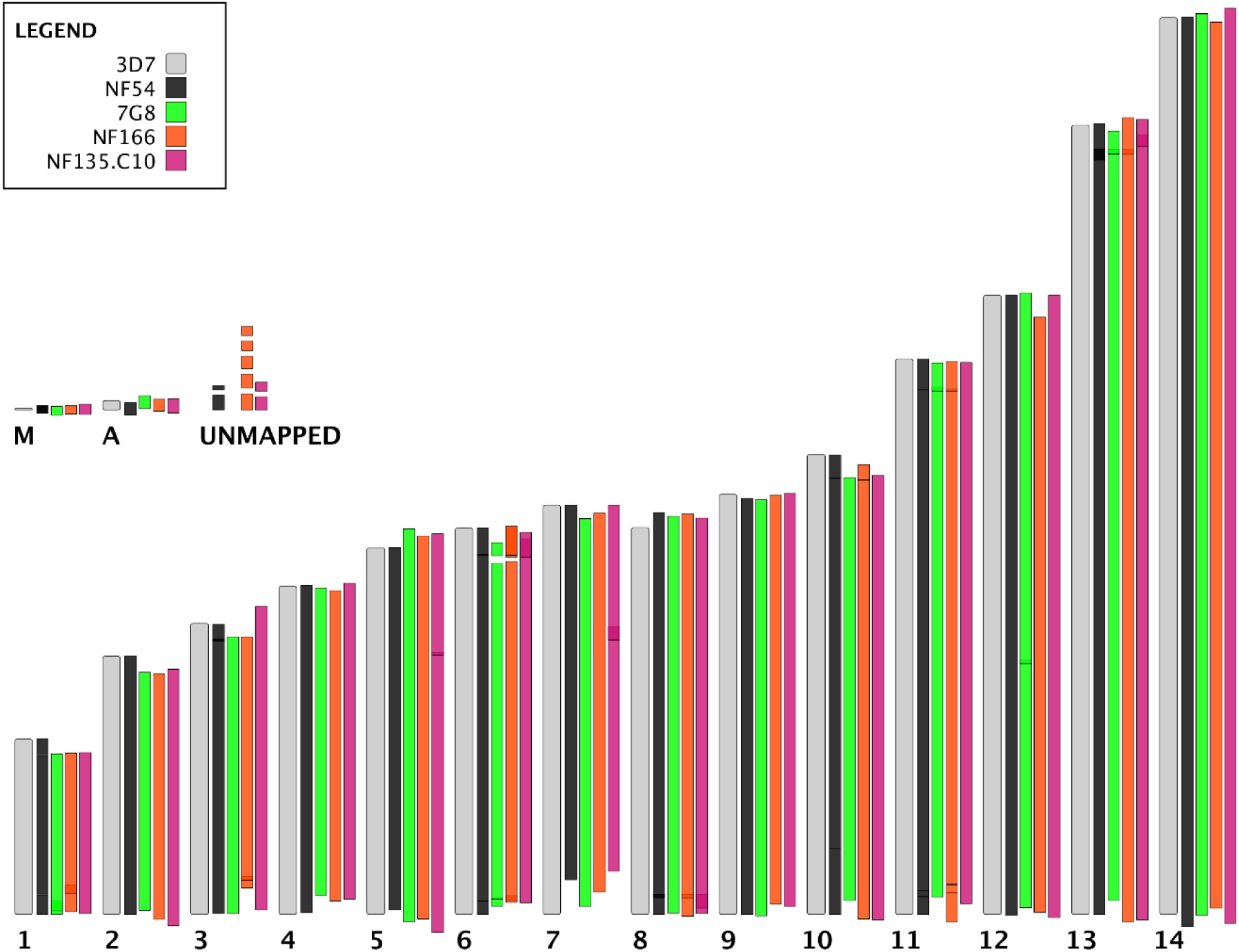
PacBio Assemblies for each PfSPZ strain reconstruct entire chromosomes in one-three continuous pieces. To determine the likely position of each non-reference contig on the 3D7 reference genome, mummer’s show-tiling program was used with relaxed settings (-g 100000 -v 50 -i 50) to align contigs to 3D7 chromosomes (top). 3D7 nuclear chromosomes (1-14) are shown in grey, arranged from smallest to largest, along with organelle genomes (M=mitochondrion, A=apicoplast). Contigs from each PfSPZ assembly (NF54: black, 7G8: green, NF166.C8: orange, NF135.C10: hot pink) are shown aligned to their best 3D7 match. A small number of contigs could not be unambiguously mapped to the 3D7 reference genome (unmapped).

**Table 1:**
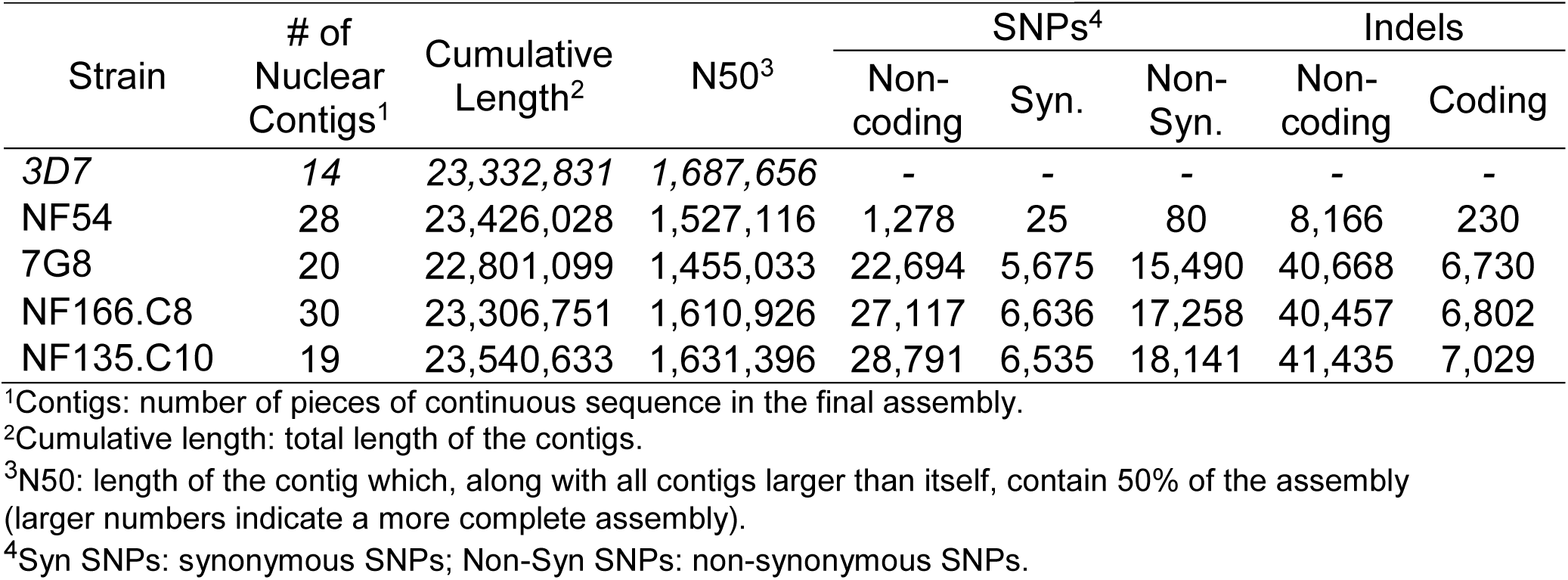
The PfSPZ strains differ from 3D7 in terms of genome size and base-pair content. Characteristics of the PacBio assembly for each strain (first four columns), with the 3D7 reference genome are shown for comparison. Single-nucleotide polymorphisms (SNPs) and indels in each PfSPZ assembly as compared to 3D7 are shown for coding and non-coding regions (last five columns).

### Structural variations in the genomes of the PfSPZ strains

Many structural variants (defined as indels or tandem repeat contractions or expansions, greater than 50 bp) were identified in each assembly by comparison to the 3D7 genome, impacting a cumulative length of 199.0 Kbp in NF166.C8 to 340.9 Kbp in NF135.C10 (**Table S5**). Many smaller variants fell into coding regions (including known pre-erythrocytic antigens), often representing variation in repeat units (**Supplemental Dataset 3**). Several larger structural variants (>10 Kbp) exist in 7G8, NF166.C8, and NF135.C10 relative to 3D7. Many of these regions contain members of multi-gene families, and as expected the number of these vary between each assembly (**Figure S2**). While all 61 3D7 *var* exon 1 sequences were present as almost identical sequences in the NF54 assembly (with the exception of small indels in several exon 1 sequences), the number of 3D7 exon 1 *var* sequences with identifiable orthologs in the three heterologous PfSPZ strains was much lower. Only a few of the identified orthologs had amino acid sequence identity higher than 75% (**Figure S2**), reflecting the exceptional sequence diversity that occurs in this gene family in and among *P. falciparum* strains. A recently characterized PfEMP1 protein expressed on the surface of NF54 sporozoites (NF54*var*^sporo^) was shown to be involved in hepatocyte invasion (Pf3D7_0809100), and antibodies to this PfEMP1 blocked hepatocyte invasion (64). No ortholog to NF54*var*^sporo^ was identified in the *var* repertoire of 7G8, NF166.C8, or NF135.C10 (**Figure S2**). It is possible that a different, strain-specific, *var* gene fulfills a similar role in each of the heterologous PfSPZ strains.

Several other large structural variants impact regions housing non-multi-gene family members, although none known to be involved in pre-erythrocytic immunity. Examples include a 31 Kbp-long tandem expansion of a region of chromosome 12 in the 7G8 assembly (also present in the previously published assembly for 7G8 (33)) and a 22.7 Kbp-long repeat expansion of a region of chromosome 5 in NF135.C10, both of which are supported by ∼200 PacBio reads. The former is a segmental duplication containing a vacuolar iron transporter (PF3D7_1223700), a putative citrate/oxoglutarate carrier protein (PF3D7_1223800), a putative 50S ribosomal protein L24 (PF3D7_1223900), GTP cyclohydrolase I (PF3D7_1224000), and three conserved *Plasmodium* proteins of unknown function (PF3D7_1223500, PF3D7_1223600, PF3D7_1224100). The expanded region in NF135.C10 represents a tandem expansion of a segment housing the gene encoding the Pf multidrug resistance protein, PfMDR1 (PF3D7_0523000), resulting in a total of four copies of this gene in NF135.C10. Other genes in this tandem expansion include those encoding an iron-sulfur assembly protein (PF3D7_0522700), a putative pre-mRNA-splicing factor DUB31 (PF3D7_0522800), a putative zinc finger protein (PF3D7_0522900) and a putative mitochondrial-processing peptidase subunit alpha protein (PF3D7_0523100). In addition, the NF135.C10 assembly contained a large translocation involving chromosomes 7 (3D7 coordinates ∼520,000 to ∼960,000) and 8 (start to coordinate ∼440,000) (**Figure S3**); the boundaries of the translocated regions contain members of multi-gene families. Independent assemblies and mapping patterns observed by aligning the PacBio reads against the NF135.C10 and 3D7 assemblies support the translocation (**Figure S4**). Furthermore, the regions of chromosomes 7 and 8 where the translocation occurs are documented recombination hotspots that were identified specifically in isolates from Cambodia, the site of origin of NF135.C10 (65).

Several structural differences in genic regions were also identified between the NF54 assembly and the 3D7 genome (**Supplemental Dataset 3**); if real, these structural variants would have important implications in the interpretation of trials using 3D7 as a homologous CHMI strain. For example, a 1,887 bp tandem expansion was identified in the NF54 assembly on chromosome 10, which overlapped the region containing liver stage antigen 1 (PfLSA-1, PF3D7_1036400). The structure of this gene in the NF54 strain was reported when PfLSA-1 was first characterized, with unique N- and C-terminal regions flanking a repetitive region consisting of several dozen repeats of a 17 amino acid motif (66, 67). The CDS of PfLSA-1 in the NF54 assembly was 5,406 bp in length but only 3,489 bp-long in the 3D7 reference. To determine if this was an assembly error in the NF54 assembly, the PfLSA-1 locus from a recently published PacBio-based assembly of 3D7 (32) was compared to that of NF54. The two sequences were identical, likely indicative of incorrect collapsing of the repeat region of PfLSA-1 in the 3D7 reference; NF54 and 3D7 PacBio-based assemblies had 79 units of the 17-mer amino acid repeat, compared to only 43 in the 3D7 reference sequence, a result further validated by the inconsistent depth of mapped Illumina reads from NF54 between the PfLSA repeat region and its flanking unique regions in the 3D7 reference (**Figure S5**). Using the same comparative rationale as above, where structural differences relative to the 3D7 reference, but common to both PabBio-based 3D7 and NF54 assemblies, are considered true, all structural variations identified in NF54 likely represent existing structural errors in the 3D7 reference genome, including 15 variants that affect genic regions (**Figure S6**). *Small sequence variants between PfSPZ strains and the reference 3D7 genome*

Very few small sequence variants were identified in NF54 compared to the 3D7 reference; 17 non-synonymous mutations were present in 15 single-copy non-pseudogene-encoding loci (**Supplemental Dataset 2**). Short indels were detected in 185 genes; many of these indels have length that is not multiple of three and occur in homopolymer runs, possibly representing remaining PacBio sequencing error. However, some may be real, as a small indel causing a frameshift in PF3D7_1417400, a putative protein-coding pseudogene that has previously been shown to accumulate premature stop codons in laboratory-adapted strains (68), and some may be of biologic importance, such as those seen in two histone-related proteins (PF3D7_0823300 and PF3D7_1020700). It has been reported that some clones of 3D7, unlike NF54, are unable to consistently produce gametocytes in long-term culture (27, 69); no polymorphisms were observed within or directly upstream of PfAP2-G (PF3D7_1222600) (**Table S6**), which has been identified as a transcriptional regulator of sexual commitment in *P. falciparum* (70). However, a non-synonymous mutation from arginine to proline (R1286P) was observed in a AP2-coincident C-terminal domain of PfAP2-L (PF3D7_0730300), a gene associated with liver stage development (71); a proline was also present at the homologous position in 7G8, NF166.C8, and NF135.C10. Two additional putative AP2 transcription factor genes (PF3D7_0613800 and PF3D7_1317200) contained a three bp deletion in NF54 relative to 3D7. Although knock-downs of PF3D7_1317200 have resulted in a loss of gametocyte production (72), neither indel was located within predicted AP2 domains, and so their biological relevance remains to be determined. 7G8, NF66.C8, and NF135.C10 also have numerous non-synonymous mutations and indels within putative AP2 genes, a small number of which fell in predicted AP2 domains. Finally, NF135.C10 has an insertion almost 200 base-pairs in length relative to 3D7 in the 3’ end of PfAP2-G; the insertion also carries a premature stop codon, leading to a considerably different C-terminal end for the transcription factor (**Figure S7**). This allele is also present in previously published assemblies for clones from Southeast Asia (33), including the culture-adapted strain DD2, and variations of this indel (without the in-frame stop codon) are also found in several non-human malaria *Plasmodium* species) (**Figure S7**), suggesting an interesting evolutionary trajectory of this sequence.

Given that no absolute correlates of protection are known for whole organism *P. falciparum* vaccines, genetic differences were assessed both across the genome and in pre-erythrocytic genes of interest in the three heterologous CHMI strains. As expected, the number of mutations between 3D7 and these three PfSPZ strains was much higher than observed for NF54, with ∼40K-55K SNPs and as many indels in each pairwise comparison, roughly randomly distributed among intergenic regions, silent and non-synonymous sites (**Table 1**, **Figure 2**), and corresponding to a pairwise SNP density relative to 3D7 of 1.9, 2.1 and 2.2 SNPs/Kbp for 7G8, NF166.C8 and NF135.C10, respectively. NF135.C10 had the highest number of unique SNPs genome-wide (SNPs not shared with other PfSPZ strains), with 5% more unique SNPs than NF166.C8 and 33% more than 7G8 (**Figure S8**). Similar trends were seen when restricting the analyses to non-synonymous SNPs (5% and 26% more than NF166.C8 and 7G8, respectively). The lower number of unique SNPs in 7G8 may be due in part to the smaller genome size of this strain. Small indels with length multiple of three (but not two) were also common in coding regions across the genome, as expected from purifying selection on mutations that disrupt the reading frame, whereas indels in non-coding regions were primarily of lengths multiple of two, from unequal crossover or replication slippage in the regions of dinucleotide repeats common in the *P. falciparum* genome (**Figure S9**), confirming what has been shown in previous work using short-read whole genome sequencing data (73).

**Figure 2:**
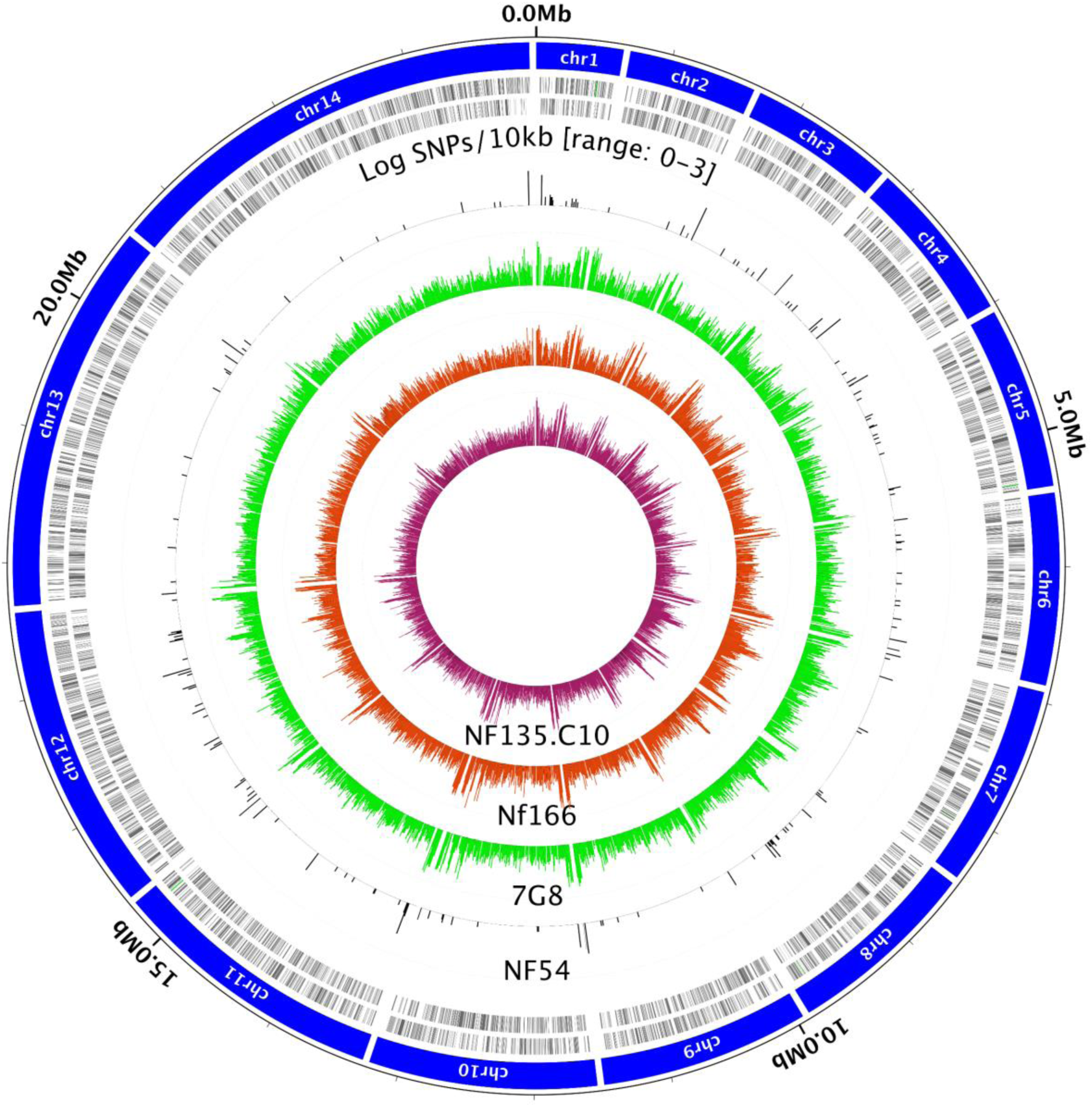
Distribution of polymorphisms in PfSPZ PacBio assemblies. Single nucleotide polymorphism (SNP) densities (log SNPs/ 10kb) are shown for each assembly; the scale [0-3] refers to the range of the log-scaled SNP density graphs - from 10^0^ to 10^3^. Inner tracks, from outside to inside, are NF54 (black), 7G8 (green), NF166.C8 (orange), and NF135.C10 (hot pink). The outermost tracks are the 3D7 reference genome nuclear chromosomes (chrm1 to chrm 14, in blue), followed by 3D7 genes on the forward and reverse strand (black tick marks).

SNPs were also common in a panel of 42 pre-erythrocytic genes known or suspected to be implicated in immunity to liver-stage parasites (see Methods; **Table S7**). While the sequence of all these loci was identical between NF54 and 3D7, there was a wide range in the number of sequence variants per locus between 3D7 and the other three PfSPZ strains, with some genes being more conserved than others. For example, PfCSP showed 8, 7 and 6 non-synonymous mutations in 7G8, NF166.C8, and NF135.C10, respectively, relative to 3D7. However, PfLSA-1 had over 100 non-synonymous mutations in all three heterologous strains relative to 3D7 (many in the repetitive region of this gene), in addition to significant length differences in the internal repeat region (**Figure S10**).

### Immunological relevance of genetic variation among PfSPZ strains

The sequence variants mentioned above may impact the ability of the immune system primed with NF54 to recognize the other PfSPZ strains, impairing vaccine efficacy against heterologous CHMI. Data from murine and non-human primate models (5,28,29,74) demonstrate that CD8^+^ T cells are required for protective efficacy; therefore, the identification of shared and unique CD8^+^ T cell epitopes across the genome in all four PfSPZ strains may help interpret the differential efficacy seen in heterologous relative to homologous CHMI. To create a comparable set of genes in which we were confident of the amino acid sequence, we subset the 5,548 protein-coding genes in the 3D7 genome annotation to 4,814 genes that (1) did not have premature stop codons, 2) were annotated with an expected number of mRNA fragments, and (3) met the above requirements in all four PfSPZ strains (in practice, this removed most, if not all, members of multi-gene families, such as *var*s, *rifin*s, *stevor*s) (**Figure 3A**). Strong-binding MHC class I epitopes in the protein sequences from these loci were identified using *in silico* epitope predictions based on HLA types common in sub-Saharan Africa populations (**Table S1**).

**Figure 3:**
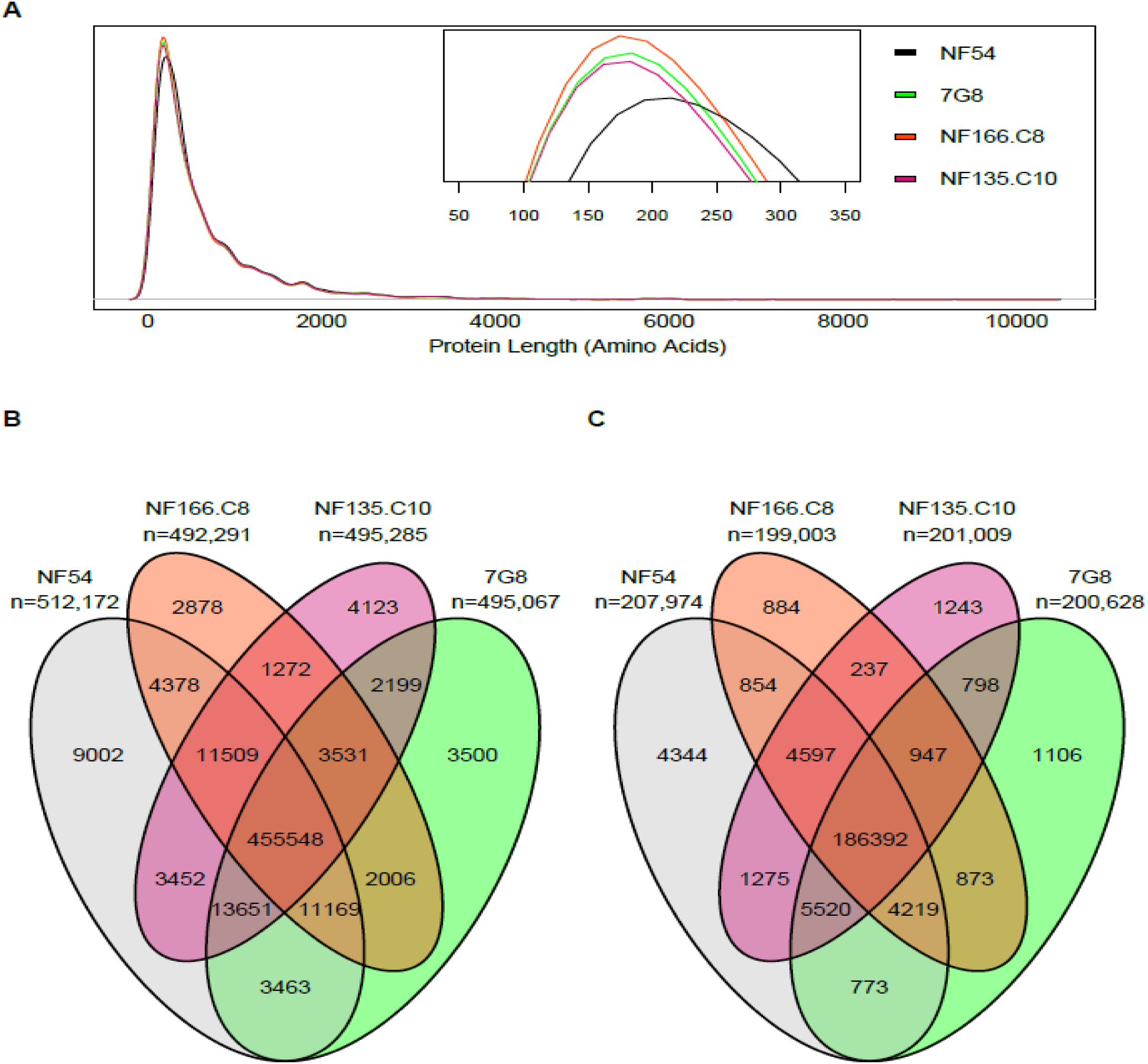
Predicted CD8^+^ T cell epitopes from amino acid sequences of the four PfSPZ strains. Using amino acid sequences from a subset of the 5,548 protein-coding genes in the 3D7 annotation (PlasmoDBv24) that were deemed to be correctly annotated between all four PfSPZ strains, CD8^+^ T cell epitopes were predicted *in silico* using netMHCpan. **A**. Density plot of amino acid sequence length distributions of 4,814 genes used for *in silico* epitope predictions. Overall distribution of gene lengths was extremely similar between all four strains, with the exception of the three heterologous CHMI strains having a slightly higher number of amino acid sequences that weren’t quite as long as they were in the NF54 genome, possibly a reflection of difficulty in mapping annotations to genetically diverse genomes. **B.** Shared and unique epitope-gene combinations in the 4,814 genes. **C**. Shared and unique epitope-gene combinations in a subset of the 4,814 genes (n=1,974) that have evidence of being expressed in the sporozoite stage of the parasite in the mosquito salivary glands.

Similar total number of epitopes (sum of unique epitopes, regardless of the HLA-type, across genes) were identified in the three heterologous CHMI strains, with NF166.C8 having the fewest (492, 291) and NF135.C10 having the most (495, 285) (**Figure 3B**). More epitopes were identified in NF54 (n=512,172) than in the three heterologous CHMI strains. This slightly higher number might be due to a more accurate mapping of the 3D7 genome annotation to the NF54 assembly compared to the other three strains (**Figure 3A**). More than 80% of epitopes detected in each strain were identical (by sequence and by locus) across all four strains, reflecting epitopes predicted in conserved regions of the 4,814 genes used in this analysis. NF54 had the highest proportion of unique epitopes (9,002, or 1.76%) (**Figure 3A**), i.e., epitopes not shared with the other PfSPZ strains, followed by NF135.C10 (4,123 or 0.83%), 7G8 (3,500 or 0.71%), and NF166.C8 (2,878, or 0.58%). After direct comparisons of NF54 to each heterologous PfSPZ strain, NF166 had the lowest proportion of epitopes not shared with NF54 (1.97%, compared to 2.24% and 2.27% for 7G8 and NF135.C10, respectively), as would follow from a positive correlation between geographic distance and genetic similarity.

To focus the analysis on genes that might be important for the protective efficacy of a whole sporozoite vaccine, the above epitope predictions were stratified by two additional gene sets: 1) 1,974 genes that were found to be expressed in the sporozoite stage of the parasite in mosquito salivary glands (54, 55), and (2) the 42 pre-erythrocytic genes of interest mentioned above. In the sporozoite-expressed gene set, similar patterns as in the genome-wide gene set above were seen, with NF135.C10 being the heterologous CHMI strain that had the highest number of epitopes not shared with any other strain (1,243, or 0.6%), and NF166 having the smallest (884, or 0.4%) (**Figure 3C**). Within the 42 pre-erythrocytic gene set, there were 117, 121, and 153 such unique epitopes present in 7G8, NF166.C8, and NF135.C10, respectively (**Figure S11**, **Table S8**). Genes where unique epitopes were identified in this gene set were also, on average, more polymorphic (**Table S7**).

Some of these variations in epitope sequences are relevant for the interpretation of the outcome of PfSPZ Vaccine trials. For example, while all four strains are identical in sequence composition in a B cell epitope potentially relevant for protection recently identified in the circumsporozoite protein (PfCSP, PF3D7_0304600) (75), another B cell epitope that partially overlaps it (76) contained an A98G amino acid difference in 7G8 and NF135.C10 relative to NF54 and NF166.C8. There was also variability in CD8^+^ T cell epitopes recognized in the Th2R region of the protein, some known for decades (77). Specifically, the PfCSP encoded by the 3D7/NF54 allele was predicted to bind to both HLA-A and HLA-C allele types, but the homologous protein segments in NF166.C8 and NF135.C10 were only recognized by HLA-A allele types, and no epitope was detected (within the HLA types studied) at that position in PfCSP encoded in 7G8 (**Figure 4**). Expansion of the analyses to additional HLA types revealed an allele (HLA-08:01) that is predicted to bind to the Th2R region of the 7G8-encoded PfCSP; however, HLA-08:01 is much more frequent in European populations (10-15%) than in African populations (1-6%) (51). Therefore, if CD8^+^ T cell epitopes in the Th2R region of 7G8 are important for protection, which is currently unknown, the level of protection against CHMI with 7G8 observed in volunteers of European descent may not be correlated with PfSPZ Vaccine efficacy in Africa. Indels and repeat regions also sometimes affected the number of predicted epitopes in each antigen; for example, an insertion in 7G8 near amino acid residue 1,600 in PfLISP-2 (PF3D7_0405300) contained additional predicted epitopes (**Figure S12**). Similar patterns in variation in epitope recognition and frequency were found in other pre-erythrocytic genes of interest, including PfLSA-3 (PF3D7_0220000), PfAMA-1 (PF3D7_1133400) and PfTRAP (PF3D7_1335900) (**Figure S12**).

**Figure 4:**
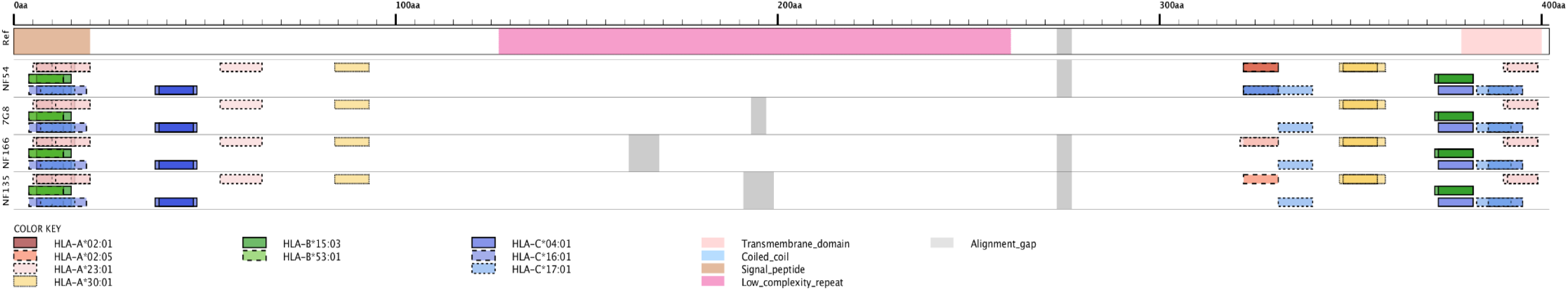
Predicted CD8^+^ T cell epitopes in the *P. falciparum* circumsporozoite protein (PfCSP). Protein domain information based on the 3D7 reference sequence of PfCSP is found in the first track, and the following tracks are epitopes predicted in the PfCSP sequences of NF54, 7G8, NF166.C8, and NF135.C10, respectively. Each box in the track is a sequence that was identified as an epitope, and the colors of each box represent the HLA type that identified the epitope.

### PfSPZ strains and global parasite diversity

The four PfSPZ strains have been adapted and kept in culture for extended periods of time. To determine if they are still representative of the malaria-endemic regions from which they were collected, we compared these strains to over 600 recent (2007-2014) clinical isolates from South America, Africa, Southeast Asia, and Oceania (**Supplemental Dataset 4**), using principal coordinates analysis (PCoA) based on SNP calls generated from Illumina whole genome sequencing data. The results confirmed the existence of global geographic differences in genetic variation previously reported (78, 79), including clustering by continent, as well as a separation of East from West Africa and of the Amazonian region from that West of the Andes (**Figure 5**). The PfSPZ strains clustered with others from their respective geographic regions, both at the genome-wide level and when restricting the data set to SNPs in the panel of 42 pre-erythrocytic antigens, despite the long-term culturing of some of these strains (**Figure 5**). An admixture analysis of South American and African clinical isolates confirmed that NF54 and NF166.C8 both have the genomic background characteristic of West Africa, while 7G8 is clearly a South American strain (**Figure S13**).

**Figure 5:**
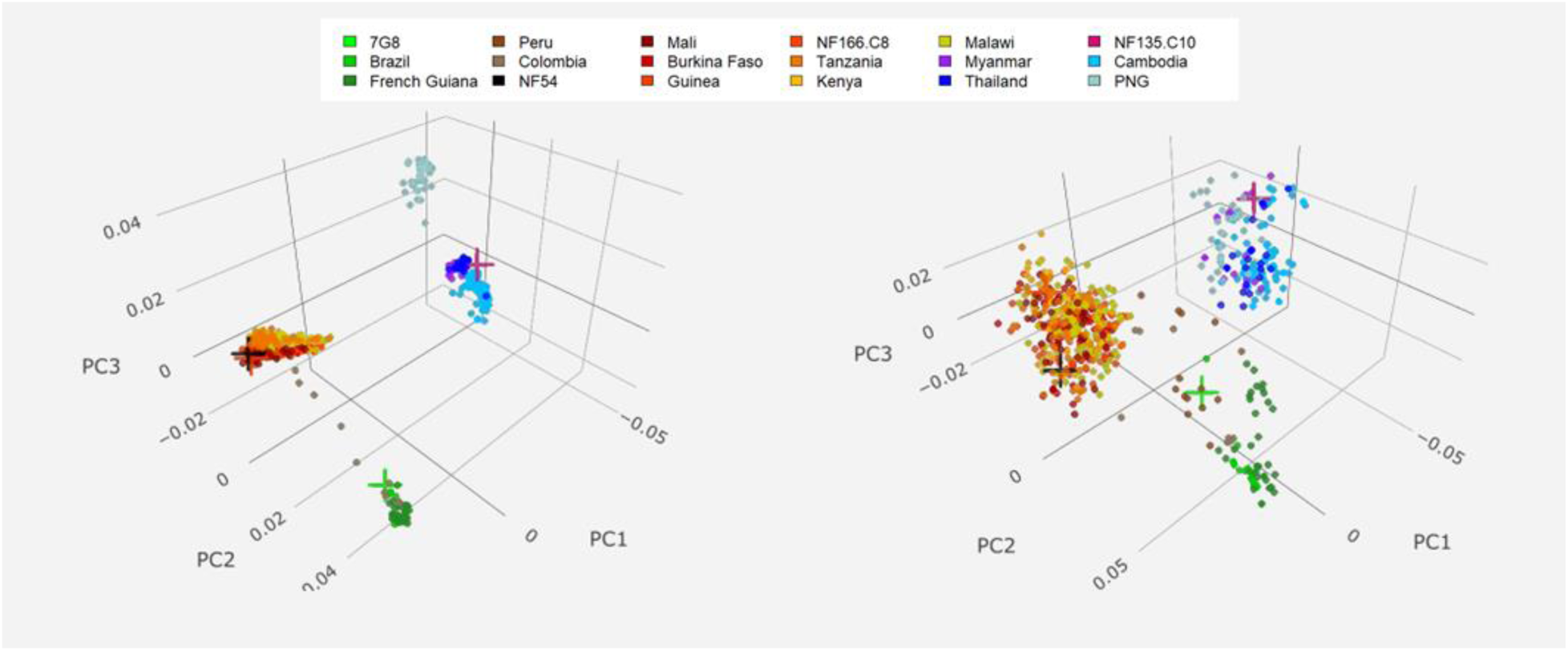
Global diversity of clinical isolates and PfSPZ strains. Principal coordinates analyses (PCoA) of clinical isolates (n=654) from malaria endemic regions and PfSPZ strains were conducted using biallelic non-synonymous SNPs across the entire genome (left, n=31,761) and in a panel of 42 pre-erythrocytic genes of interest (right, n=1,060). For the genome-wide dataset, coordinate 1 separated South American and African isolates from Southeast Asian and Papua New Guinean isolates (27.6% of variation explained), coordinate 2 separated African isolates from South American isolates (10.7%), and coordinate 3 separated Southeast Asian isolates from Papua New Guinea (PNG) isolates (3.0%). Similar trends were found for the first two coordinates seen for the pre-erythrocytic gene data set (27.1 and 12.6%, respectively), but coordinate 3 separated isolates from all three regions (3.8%). In both datasets, NF54 (black cross) and NF166.C8 (orange cross) cluster with West African isolates (isolates labeled in red and dark orange colors), 7G8 (bright green cross) cluster with isolates from South America (greens and browns), and NF135.C10 (hot pink cross) clusters with isolates from Southeast Asia (purples and blues).

NF135.C10 was isolated in the early 1990s (14), at a time when resistance to chloroquine and sulfadoxine-pyrimethamine resistance was entrenched and resistance to mefloquine was emerging (80, 81), and carries signals from this period of drug pressure. Four copies of MDR-1 were identified in NF135.C10 (**Table S9**); however, two of these copies appeared to have premature stop codons introduced by SNPs and/or indels, leaving potentially only two functional copies in the genome. While NF135.C10 also had numerous point mutations relative to 3D7 in genes such as PfCRT (conveying chloroquine resistance), and PfDHPS and PfDHR (conveying sulfadoxine-pyrimethamine resistance), NF135.C10 was isolated before the widespread deployment of artemisinin-based combination therapies (ACTs), and had the wild-type allele in the locus that encodes the kelch protein in chromosome 13 (PfK13), with no mutations known to convey artemisinin resistance detected in the propeller region (**Table S10**).

The emergence in Southeast Asia of resistance to antimalarial drugs, including artemisinins and drugs used in artemisinin-based combination treatments, is thought to underlie the complex and dynamic parasite population structure in the region (82). Several relatively homogeneous subpopulations, whose origin is likely linked to the emergence and rapid spread of drug resistance mutations, exist in parallel with a sensitive subpopulation that reflects the ancestral population in the region (referred to as KH1), and another subpopulation of admixed genomic background (referred to as KHA), possibly the source of the drug-resistant subpopulations or the result of a secondary mix of resistant subpopulations (61,62,83,84). This has been accompanied by reports of individual K13 mutations conferring artemisinin resistance occurring independently on multiple genomic backgrounds (85). To determine the subpopulation to which NF135.C10 belongs, an admixture analysis was conducted using isolates from Southeast Asia and Oceania, including NF135.C10. Eleven total populations were detected, of which seven contained Cambodian isolates (**Figure 6**). Both admixture and hierarchical clustering analyses suggest that NF135.C10 is representative of the previously described admixed KHA subpopulation (61, 62) (**Figure 6**), implying that NF135.C10 is representative of a long-standing admixed population of parasites in Cambodia rather than one of several subpopulations thought to have arisen more recently following the emergence of drug resistance.

**Figure 6:**
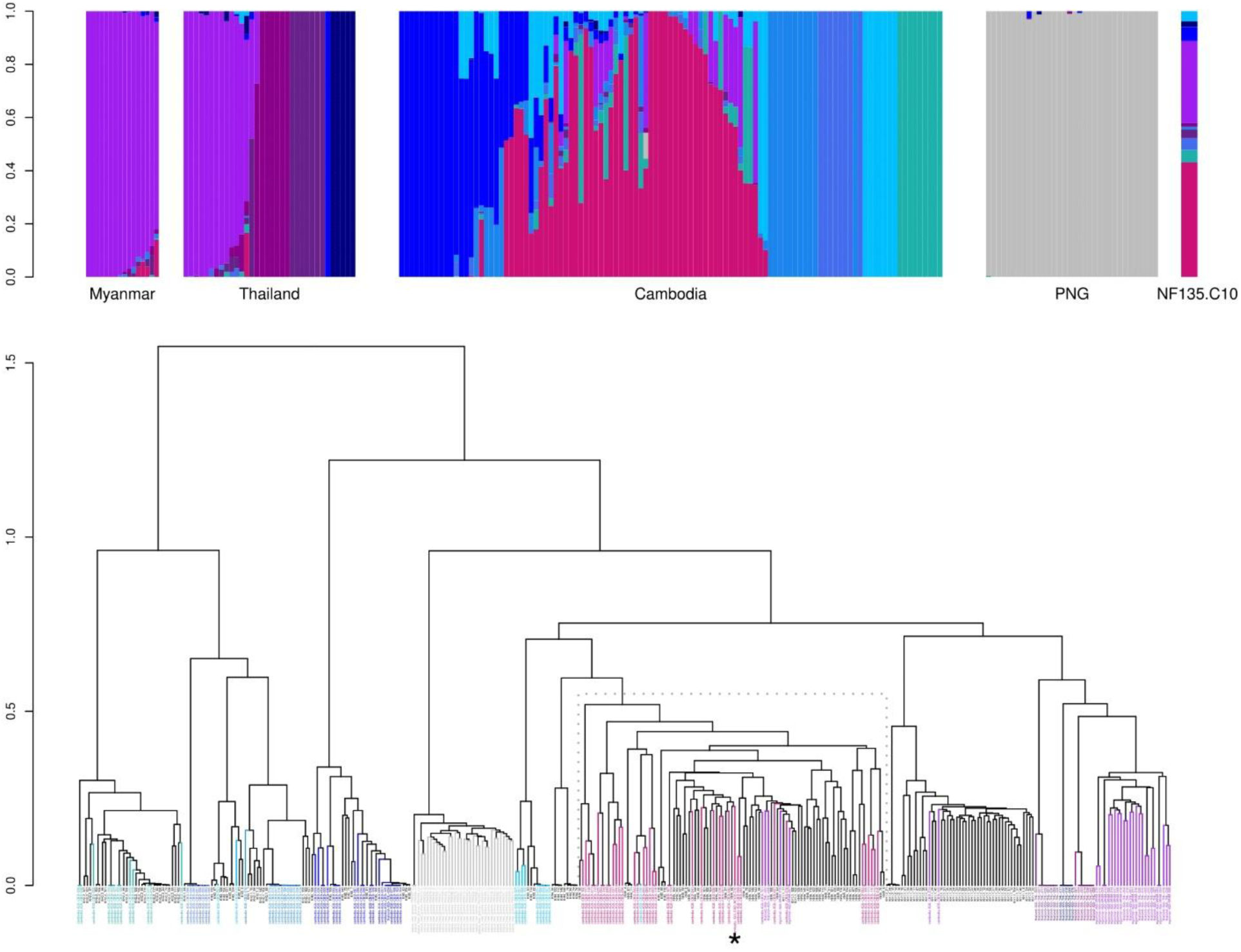
NF135.C10 is part of an admixed population of clinical isolates from Southeast Asia. Top: admixture plots for clinical isolates from Myanmar (n=16), Thailand (n=34), Cambodia (n=109), Papua New Guinea (PNG, n=34) and NF135.C10 are shown. Each sample is a column, and the height of the different colors in each column corresponds to the proportion of the genome assigned to each *K* population by the model. Bottom: hierarchical clustering of the Southeast Asian isolates used in the admixture analysis (n=193; colored by their assigned subpopulation) and previously characterized Cambodian isolates (n=167; black; labels correspond to clusters identified in reference 61) place NF135.C10 (star) with samples from the previously identified KHA admixed population (shown in grey dashed box). Each node ends with a sample, and the y-axis represents distance between clusters.

## Discussion

Whole organism sporozoite vaccines have provided variable levels of protection in initial clinical trials; the radiation-attenuated PfSPZ Vaccine has been shown to protect >90% of subjects against homologous CHMI at 3 weeks after the last dose in 5 clinical trials in the U.S. (6, 8) and Germany (11). However, efficacy has been lower against heterologous CHMI (8, 9), and in field studies in a region of intense transmission, in Mali, at 24 weeks (10). Interestingly, for the exact same immunization regimen, protective efficacy by proportional analysis was greater in the field trial in Mali (29%) than it was against heterologous CHMI with Pf 7G8 in the US at 24 weeks after last dose of vaccine (8%) (8, 10). While evidence shows that whole organism-based vaccine efficacy can be improved by adjusting the vaccine dose and schedule (11), further optimization of such vaccines will be facilitated by a thorough understanding of the genotypic and immunologic differences among the PfSPZ strains and between them and parasites in malaria endemic regions.

A recent study examined whole genome short-read sequencing data to characterize NF166.C8 and NF135.C10 through SNP calls, and identified a number of non-synonymous mutations at a few loci potentially important for the efficacy of CPS, the foundation for PfSPZ-CVac (17). The analyses described here, using high-quality *de novo* genome assemblies, expand the analysis to hard-to-call regions, such as those containing gene families, repeats and other low complexity sequences. The added sensitivity enabled the thorough genomic characterization of these and additional vaccine-related strains, and revealed a considerably higher number of sequence variants than can be called using short read data alone, as well as indels and structural variants between assemblies. For example, the insertion close to the 3’ end of PfAP2-G detected in NF135.C10 and shared by DD2 has not, to the best of our knowledge, been reported before, despite the multiple studies highlighting the importance of this gene in sexual commitment in *P. falciparum* strains, including DD2 (70). Long-read sequencing also confirmed that differences observed between NF54 and 3D7 in a major liver stage antigen, PfLSA-1, result from one of a small number of errors lingering in the reference 3D7 genome, which is being continually updated and improved (34). Confirmation that NF54 and 3D7 are identical at this locus is critical when 3D7 has been used as a homologous CHMI in whole sporozoite, NF54-based vaccine studies. Furthermore, the comprehensive sequence characterization of variant surface antigen-encoding loci, such as PfEMP1-encoding genes, will enable the use of the PfSPZ strains to study the role of these protein families in virulence, naturally acquired immunity and vaccine-induced protection (86).

The comprehensive genetic and genomic studies reported herein were designed to provide insight into the outcome of homologous and heterologous CHMI studies, and to determine whether the CHMI strains can be used as a proxy for strains present in the field. Comparison of genome assemblies confirmed that NF54 and 3D7 have remained genetically very similar over time, and that 3D7 is an appropriate homologous CHMI strain. As expected, 7G8, NF166.C8, and NF135.C10 were genetically very distinct from NF54 and 3D7, with thousands of differences across the genome including dozens in known pre-erythrocytic antigens. The identification of sequence variants (both SNPs and indels) within transcriptional regulators, such as the AP2 family, may assist in the study of different growth phenotypes in these strains. NF166.C8 and NF135.C10 merozoites enter the blood stream several days earlier than those of NF54 (15), suggesting that NF54 may develop more slowly in hepatocytes than do the other two strains. Therefore, mutations in genes associated with liver-stage development (as was observed with PfAP2-L) may be of interest to explore further. Finally, comparison of the PfSPZ strains to whole genome sequencing data from clinical isolates shows that, at the whole genome level, they are indeed representative of their geographical regions of origin.

These results can assist in the interpretation of CHMI studies in multiple ways. First, in the 42 pre-erythrocytic genes of interest, the Brazilian strain 7G8, one of the first strains to be used in heterologous CHMI studies, and the Guinean strain NF166.C8 share a similar number of epitopes with NF54. Furthermore, both NF166.C8 and NF135.C10 encode more unique non-synonymous SNPs genome-wide (19% and 33%, respectively) and epitopes in these genes relative to NF54 than does 7G8. These results show that the practice of equating geographic distance to genetic differentiation is not always valid, and that the interpretation of CHMI studies should rest upon thorough genome-wide comparisons. Lastly, of all PfSPZ strains, NF135.C10 is the most genetically distinct from NF54, including in point mutations, unique epitopes, and its placement within the global parasite population. We do not know whether the peptide sequences of the actual epitope targets of vaccine-induced protective immunity are equally distinct in NF135.C10. However, if proteome-wide genetic divergence is the primary determinant of differences in protection against different parasites, the extent to which NF54-based immunization protects against CHMI with NF135.C10 is important in understanding the ability of PfSPZ Vaccine and other whole-organism malaria vaccines to protect against diverse parasites present world-wide. These conclusions are drawn from genome-wide analyses and from subsets of genes for which a role in whole-sporozoite-induced protection is suspected but not experimentally established. Conclusive statements regarding cross-protection will require the additional knowledge of the genetic basis of whole-organism vaccine protection.

Without more information on the epitope targets of protective immunity induced by PfSPZ vaccines, it is difficult to rationally design multi-strain PfSPZ vaccines. However, these data and analysis approach has the potential to be applied to rationally design multi-strain sporozoite-based vaccines once knowledge of those critical epitope sequences is available, and characterization of a variety of *P. falciparum* strains may facilitate the development of region-specific or multi-strain vaccines with greater protective efficacy. Support for a genomics-guided approach to guide such next-generation vaccines can be found in other whole organism parasitic vaccines. Field trials testing the efficacy of first-generation whole killed-parasite vaccines against *Leishmania* had highly variable results (87). While most studies failed to show protection, indicating that killed, whole-cell vaccines for leishmaniasis may not produce the necessary protective response, a trial demonstrating significant protection utilized a multi-strain vaccine, with strains collected from the immediate area of the trial (88), highlighting the importance of understanding the distribution of genetic diversity in pathogen populations. In addition, a highly efficacious non-attenuated, three-strain, whole organism vaccine exists against *Theileria parva*, a protozoan parasite that causes East coast fever in cattle. This vaccine, named Muguga Cocktail, consists of a mix of three live strains of *T. parva* that are administered in an infection-and-treatment method, similar to the approach utilized by PfSPZ-CVac. It has been shown recently that two of the strains are genetically very similar, possibly clones of the same isolates (89). Despite this, the vaccine remains highly efficacious and in high demand (90). In addition, the third vaccine strain in the Muguga Cocktail is quite distinct from the other two, with ∼5 SNPs/kb (89), or about twice the SNP density seen between NF54 and other PfSPZ strains. These observations suggest that an efficacious multi-strain vaccine against a highly variable parasite species does not need to be contain a large number of strains, but that the inclusion of highly divergent strains may be warranted. These results also speak to the promise of multi-strain vaccines against highly diverse pathogens, including apicomplexans with large genomes and complex life cycles.

Next-generation whole genome sequencing technology has opened many avenues for infectious disease research and holds great promise for informing vaccine design. While most malaria vaccine development has occurred before the implementation of regular use of whole genome sequencing, the tools now available allow the precise characterization and informed selection of vaccine strains early in the development process. These results will greatly assist in the interpretation of clinical trials using these strains for CHMI.

## Declarations

### Ethics approval and consent to participate

Not applicable

### Consent for publication

Not applicable

### Availability of data and material

Raw sequencing data and assemblies generated for this manuscript have been uploaded to NCBI. See Dataset 1 for BioSample identification numbers for assemblies and whole genome sequence data for all samples used in these analyses, including publicly available data.

### Competing interests

T.L., B.K.L.S., and S.L.H. are salaried employees of Sanaria Inc., the developer and owner of PfSPZ Vaccine, and the supplier of the PfSPZ strains sequenced in this study. In addition, S.L.H. and B.K.L.S. have a financial interest in Sanaria Inc. All other authors declare no competing financial interests.

### Funding

National Institutes of Health (NIH) awards U19 AI110820 and R01 AI141900, and a “Dean’s Challenge Award”, an intramural program from the University of Maryland School of Medicine, provided funds to sequence the PfSPZ strains and Pf clinical isolates, and support for K.A.M., E.F.D., A.D., M.A., A.O., K.L., L.S., L.T., M.T., S.T.H., C.M.F., C.V.P. and J.C.S.. A.O. and C.V.P. were also supported by the Howard Hughes Medical Institute. The parasites provided by Sanaria were produced in part by funding from two SBIR grants from National Institute of Allergy and Infectious Diseases, NIH, 2R44AI058375 and 5R44AI055229-09A1. Collection of clinical isolates that were sequenced for this study were funded by U19 AI089683, grants from CNPq (590106/2011-2), grants from the Armed Forces Health Surveillance Center/Global Emerging Infections Surveillance and Response System (WR1576 and WR2017), the NIH International Centers of Excellence in Malaria Research program (U19 AI089681) (Brazil), and the Howard Hughes Medical Institute. S.K. and A.M.P. were supported by the Intramural Research Program of the National Human Genome Research Institute, NIH (1ZIAHG200398). P.T.R. was supported by a scholarship from the Conselho Nacional de Desenvolvimento Científico e Tecnológico (CNPq), which also provided a senior researcher scholarship to M.U.F.

### Authors’ contributions

K.A.M. designed the study, performed analyses, and wrote the paper. J.C.S., C.V.P., and S.L.H. designed the study, contributed samples and funding, and wrote the paper. E.F.D., A.D., J.C., E.M.S., A.D., S.K., A.M.P, and A.O. contributed analyses. Z.S., M.A., T.L., L.S., L.T. preformed sample preparation and sequencing. P.T.R., K.E.L., M.D.S., K.J., C.L., D.L.S., M.U.F., M.N., M.K.L., M.T., R.W.S., S.T.H., C.M.F, and B.K.L.S contributed samples, funding, and wrote the paper.

## Acknowledgements

We would like to thank the study teams that assisted in sample collections in Mali, Malawi, Cambodia, Thailand, and Myanmar. We would like to acknowledge Drs. Terrie Taylor and Don Mathanga for their assistance with collection of clinical isolates in Malawi.

## Authors’ information (optional)

Additional author information for M. D. S, K. J., C. L., D. L. S (affiliated with the Armed Forces Research Institute of Medical Sciences): Material has been reviewed by the Walter Reed Army Institute of Research. There is no objection to its presentation and/or publication. The opinions or assertions contained herein are the private views of the author, and are not to be construed as official, or as reflecting true views of the Department of the Army or the Department of Defense. The investigators have adhered to the policies for protection of human subjects as prescribed in AR 70–25.

