## Supplemental Tables and Figures for "Strains used in whole organism *Plasmodium falciparum* vaccine trials differ in genome structure, sequence, and immunogenic potential"

**Table S1.** Frequency of HLA types for MHC class-1 epitopes in West and East Africa

| | HLA Type - MHC Class 1[1] | | | | | | | | |
|---|---|---|---|---|---|---|---|---|---|
| | HLA-A02:01 | HLA-A02:05 | HLA-A23:01 | HLA-A30:01 | HLA-B15:03 | HLA-B53:01 | HLA-C04:01 | HLA-C16:01 | HLA-C17:01 |
| Mali | 0.083 | | 0.228 | 0.141 | 0.069 | 0.159 | 0.213 | 0.283 | 0.143 |
| BF (Rimabaibe) | 0.138 | | | | 0.138 | | | 0.16 | 0.106 |
| BF (Mossi) | | | | | | 0.208 | | 0.179 | 0.151 |
| BF (Fulani) | | 0.092 | | | | 0.082 | | 0.214 | 0.02 |
| Equatorial Guinea | | | | | | | | 0.105 | |
| Tanzania | | | | | 0.143 | 0.091 | | | |
| Kenya (Luo) | 0.115 | 0.087 | 0.089 | 0.084 | 0.089 | 0.068 | 0.132 | 0.045 | 0.087 |
| Kenya (Nandi) | 0.118 | | | | 0.079 | 0.088 | 0.115 | 0.044 | 0.102 |

[1]Allele frequencies queried from the Allele Frequency database (main manuscript reference 51, accessed October 2016 and from the literature (main manuscript references 52 and 53). BF: Burkina Faso.

**Table S2.** Pacific Biosciences (PacBio) long-read (P6-P4 Chemistry) whole genome sequencing data

| Strain | Average Library Size (bp) | # of SMRT Cells | Total # of Reads | Total # of bp | Mean Read Length (bp) | Max Read Length (bp) | Coverage[1] |
|---|---|---|---|---|---|---|---|
| NF54 | 18,800 | 4 | 557,996 | 4,146,771,099 | 7,432 | 51,499 | 151.6 |
| NF166.C8 | 19,749 | 4 | 768,860 | 5,267,844,194 | 6,856 | 49,121 | 194.5 |
| 7G8 | 19,490 | 4 | 292,390 | 2,558,840,554 | 8,754 | 55,366 | 99.1 |
| NF135.C10 | 17,035 | 4 | 357,521 | 2,768,837,384 | 7,757 | 50,024 | 106.9 |

[1]Calculated by the average read length multiplied by number of reads that mapped to the 3D7 reference (PlasmoDBv24), divided by 3D7 genome size

**Table S3.** Illumina short-read whole genome sequencing data (HiSeq 2500)

| Strain | Average Library Size (bp) | Read Length (bp) | Total # of Reads | Total # of bp | Coverage[1] |
|---|---|---|---|---|---|
| NF54 | 403 | 100 | 39,384,560 | 3,977,840,560 | 169 |
| NF166.C8 | 359 | 150 | 58,890,306 | 8,892,436,206 | 263 |
| 7G8 | 322 | 150 | 54,894,060 | 8,289,003,060 | 340 |
| NF135.C10 | 330 | 150 | 56,189,696 | 8,484,644,096 | 268 |

[1]Calculated by the average read length multiplied by number of reads that mapped to the 3D7 reference (PlasmoDBv24), divided by 3D7 genome size

**Table S4.** Comparison of 7G8 assembly generated for this manuscript (mns.) to a previously generated 7G8 assembly available on PlasmoDB (v41). Variants were identified by comparing each assembly to the 3D7 reference genome using nucmer's show-snps function (see Methods). Nm3=non-multiple of 3; m3=multiple of 3.

| Genomic Partition | Type of Variant | Mns. | PlasmoDB |
|---|---|---|---|
| Coding | deletion.Nm3 | 1,088 | 1,171 |
| | deletion.m3 | 2,391 | 2,369 |
| | insertion.Nm3 | 1,125 | 1,040 |
| | insertion.m3 | 2,143 | 2,150 |
| | point mutation | 21,452 | 21,165 |
| Non-coding | deletion.Nm3 | 16,094 | 17,411 |
| | deletion.m3 | 3,616 | 3,703 |
| | insertion.Nm3 | 17,160 | 16,009 |
| | insertion.m3 | 3,616 | 3,545 |
| | point mutation | 22,907 | 22,694 |

**Table S5.** Structural variants (>50 bps) in each PfSPZ assembly as compared to the 3D7 reference genome[1], including number and cumulative length in base pairs.

| Variant Type | NF54 | | 7G8 | | NF166.C8 | | NF135.C10 | |
|---|---|---|---|---|---|---|---|---|
| | number | bp | number | bp | number | bp | number | bp |
| Insertions | 4 | 476 | 170 | 34,060 | 161 | 16,782 | 180 | 31,721 |
| Deletions | 2 | 131 | 187 | 16,030 | 156 | 22,860 | 209 | 21,654 |
| Tandem_expansion | 26 | 23,672 | 102 | 61,958 | 114 | 95,085 | 118 | 162,337 |
| Tandem_contraction | 5 | 1,416 | 137 | 22,077 | 110 | 31,885 | 135 | 20,872 |
| Repeat_expansion | 2 | 738 | 19 | 12,500 | 22 | 14,070 | 23 | 20,526 |
| Repeat_contraction | 0 | 0 | 21 | 54,963 | 15 | 18,364 | 27 | 83,759 |
| **Total** | **39** | **26,433** | **636** | **201,318** | **578** | **199,046** | **692** | **340,869** |

[1]A small number of structural variants detected here may be due to existing errors in the 3D7 reference genome (see Figure S2)

**Table S6.** Variants identified in loci encoding predicted AP2 transcription factors

| Gene ID | Product Description | NF54 | | | 7G8 | | | NF166.C8 | | | NF135.C10 | | | Mutations occur in AP2 domains? |
|---|---|---|---|---|---|---|---|---|---|---|---|---|---|---|
| | | SNPs | | AA Length Difference | SNPs | | AA Length Difference | SNPs | | AA Length Difference | SNPs | | AA Length Difference | |
| | | S | NS | | S | NS | | S | NS | | S | NS | | |
| PF3D7_0404100 | AP2 domain transcription factor AP2-SP2, putative | 0 | 0 | 0 | 8 | 6 | -14 | 10 | 5 | -21 | 1 | 6 | -52 | no |
| PF3D7_0420300 | AP2 domain transcription factor, putative | 0 | 0 | 0 | 4 | 5 | 15 | 5 | 7 | 18 | 4 | 7 | -21 | 7G8[3274:H->Y] |
| PF3D7_0516800 | AP2 domain transcription factor AP2-O2, putative | 0 | 0 | 0 | 2 | 6 | 52 | 2.5 | 7.5 | 80 | 2.5 | 9.5 | 62 | no |
| PF3D7_0604100 | AP2 domain transcription factor | 0 | 0 | 0 | 0 | 1 | 13 | 0 | 1 | -3 | 0 | 1 | -2 | no |
| PF3D7_0611200 | AP2 domain transcription factor, putative | 0 | 0 | 0 | 1 | 0 | 0 | 0 | 0 | 0 | 0 | 0 | 0 | no |
| PF3D7_0613800 | AP2 domain transcription factor, putative | 0 | 0 | -1 | 9.5 | 28.5 | -9 | 12.5 | 14.5 | -5 | 9.5 | 25.5 | 24 | 7G8, NF166, NF135.C10[3713:N->D] |
| PF3D7_0622900 | AP2 domain transcription factor AP2Tel | 0 | 0 | 0 | 0 | 0 | 10 | 0 | 0 | -6 | 0 | 0 | -2 | no |
| PF3D7_0730300 | AP2 domain transcription factor AP2-L | 0 | 1 | 0 | 1 | 3 | 8 | 1 | 3 | 2 | 2 | 4 | 13 | no |
| PF3D7_0802100 | AP2 domain transcription factor, putative | 0 | 0 | 0 | 5 | 6 | 10 | 2 | 3 | -6 | 2 | 2 | -8 | no |
| PF3D7_0934400 | AP2 domain transcription factor, putative | 0 | 0 | 0 | 0 | 0 | 0 | 0 | 1 | 0 | 0 | 0 | 0 | no |
| PF3D7_1007700 | AP2 domain transcription factor, putative, AP2-I | 0 | 0 | 0 | 0 | 0 | 19 | 0 | 1 | 27 | 1.5 | 4.5 | 37 | no |
| PF3D7_1107800 | AP2 domain transcription factor, putative | 0 | 0 | 0 | 0 | 1 | -5 | 1 | 0 | -7 | 0 | 2 | 1 | no |
| PF3D7_1115500 | AP2 domain transcription factor, putative | 0 | 0 | 0 | 0 | 0 | 0 | 0 | 0 | 0 | 0 | 0 | 0 | no |
| PF3D7_1139300 | AP2 domain transcription factor, putative | 0 | 0 | 0 | 1 | 7 | -9 | 1 | 5 | -4 | 1 | 6 | -5 | no |
| PF3D7_1143100 | AP2 domain transcription factor AP2-O, putative | 0 | 0 | 0 | 0 | 3 | -1 | 0 | 2 | -21 | 0 | 3 | -29 | no |
| PF3D7_1222400 | AP2 domain transcription factor | 0 | 0 | 0 | 3 | 2 | 11 | 3 | 2 | 9 | 1 | 3 | -13 | no |
| PF3D7_1222600 | AP2 domain transcription factor AP2-G | 0 | 0 | 0 | 1 | 7 | 1 | 1 | 7 | 2 | 1 | 7 | 70 | no |
| PF3D7_1239200 | AP2 domain transcription factor, putative | 0 | 0 | 0 | 0 | 0 | 5 | 0 | 1 | 1 | 0 | 1 | 1 | no |

| | | | | | | | | | | | | | | |
|---|---|---|---|---|---|---|---|---|---|---|---|---|---|---|
| PF3D7_1305200 | AP2 domain transcription factor, putative | 0 | 0 | 0 | 0 | 0 | 0 | 0 | 0 | 0 | 0 | 0 | 0 | no |
| PF3D7_1317200 | AP2 domain transcription factor, putative | 0 | 0 | -1 | 0 | 0 | -22 | 0 | 0 | -37 | 0 | 0 | -38 | no |
| PF3D7_1342900 | AP2 domain transcription factor, putative | 0 | 0 | 0 | 1 | 4 | 12 | 3 | 9 | -19 | 1 | 5 | -69 | no |
| PF3D7_1350900 | AP2 domain transcription factor AP2-O4, putative | 0 | 0 | 0 | 1 | 8 | -4 | 2 | 8 | -5 | 3 | 8 | -5 | - |
| PF3D7_1408200 | AP2 domain transcription factor AP2-G2, putative | 0 | 0 | 0 | 1 | 8 | -8 | 2 | 8 | -16 | 3 | 8 | -39 | no |
| PF3D7_1429200 | AP2 domain transcription factor AP2-O3, putative | 0 | 0 | 0 | 0 | 0 | 18 | 0 | 0 | 6 | 0 | 0 | -6 | no |
| PF3D7_1449500 | AP2 domain transcription factor, putative | 0 | 0 | 0 | 2 | 1 | 2 | 1 | 1 | -1 | 1 | 1 | 4 | no |
| PF3D7_1456000 | AP2 domain transcription factor, putative | 0 | 0 | 0 | 0 | 2 | -11 | 1 | 4 | 16 | 3 | 2 | -16 | no |
| PF3D7_1466400 | AP2 domain transcription factor AP2-SP | 0 | 0 | 0 | 4 | 5 | 0 | 5 | 7 | 1 | 4 | 7 | 0 | no |

**Table S7.** Variants identified in potentially important pre-erythrocytic loci in 7G8, NF166.C8, and NF135.C10

| Gene ID | Product Description | Gene Panel[1] | | | 7G8 | | NF166.C8 | | NF135.C10 | |
|---|---|---|---|---|---|---|---|---|---|---|
| | | Pre-erythroctyic Antigens | PfSPZ Vaccinees | PfSPZ-CVac Vaccinees | Synonymous (S) and Non-synonymous (NS) Variants | | | | | |
| | | | | | S | NS | S | NS | S | NS |
| PF3D7_0206900.2 | merozoite surface protein 5 (MSP5) | x | x | | 0 | 1 | 1 | 0 | 0 | 0 |
| PF3D7_0207000 | merozoite surface protein 4 (MSP4) | x | | x | 0 | 3 | 0 | 0 | 0 | 0 |
| PF3D7_0220000 | liver stage antigen 3 (LSA3) | x | | | 12 | 42 | 13 | 29 | 12 | 62 |
| PF3D7_0304600 | circumsporozoite protein (CSP) | x | x | x | 33 | 8 | 4 | 7 | 2 | 6 |
| PF3D7_0312400 | pfGSK3 | | x | | 0 | 0 | 0 | 0 | 0 | 1 |
| PF3D7_0405300 | liver specific protein 2 (LISP2) | x | | x | 10 | 33 | 11 | 29 | 3 | 21 |
| PF3D7_0408600 | sporozoite invasion-associated protein 1 (SIAP1) | x | | | 0 | 4 | 0 | 3 | 0 | 5 |
| PF3D7_0408700 | perforin-like protein 1 (PLP1/SPECT2) | x | | | 1 | 1 | 1 | 2 | 0 | 2 |
| PF3D7_0420000 | Zinc finger protein, putative | | | x | 7 | 19 | 7 | 11 | 7 | 16 |
| PF3D7_0522400 | Conserved Plasmodium protein, unknown function | | | x | 0 | 5 | 3 | 8 | 5 | 10 |
| PF3D7_0725100 | Conserved Plasmodium membrane protein, unknown function | | | x | 1 | 1 | 1 | 0 | 1 | 0 |
| PF3D7_0726400 | Conserved Plasmodium membrane protein, unknown function | | | x | 2 | 9 | 4 | 7 | 2 | 8 |
| PF3D7_0728600 | zinc finger, C3HC4 type, putative | | x | | 1 | 2 | 4 | 2 | 3 | 5 |
| PF3D7_0812300 | sporozoite surface protein 3 (SSP3) | x | | | 0 | 0 | 0 | 1 | 0 | 1 |
| PF3D7_0815500 | Conserved Plasmodium protein, unknown function | | | x | 0 | 0 | 0 | 0 | 0 | 0 |
| PF3D7_0826100 | E3 ubiquitin-protein ligase, putative | | | x | 4 | 7 | 3 | 8 | 10 | 15 |
| PF3D7_0828100 | conserved Plasmodium protein, unkown function | | x | | 1 | 2 | 1 | 1 | 2 | 2 |
| PF3D7_0830300 | sporozoite invasion-associated protein-2 (SIAP-2) | x | | | 0 | 4 | 0 | 4 | 0 | 5 |

| | | | | | | | | | | |
|---|---|---|---|---|---|---|---|---|---|---|
| PF3D7_0906700 | Leucine-rich repeat protein (PfLR9) | | x | | 0 | 0 | 0 | 0 | 0 | 0 |
| PF3D7_1021700 | Conserved Plasmodium membrane protein, unknown function | | | x | 3 | 12 | 3 | 6 | 4 | 10 |
| PF3D7_1030200 | claudin-like apicomplexan microneme protein, putative | | x | | 2 | 1 | 1 | 0 | 1 | 0 |
| PF3D7_1035300 | Glutamate-rich protein (GLURP) | | | x | 4 | 18 | 1 | 2 | 6 | 28 |
| PF3D7_1036400* | Liver stage antigen 1 (LSA1) | x | | x | > 100 | > 100 | > 100 | 89 | > 100 | > 100 |
| PF3D7_1121600 | exported protein 1 (EXP1) | x | | | 0 | 1 | 0 | 1 | 0 | 2 |
| PF3D7_1133400 | apical membrane antigen 1 (AMA1) | x | | | 1 | 31 | 3 | 30 | 3 | 31 |
| PF3D7_1138400 | Guanylyl cyclase (GCalpha) | | | x | 1 | 3 | 1 | 5 | 2 | 7 |
| PF3D7_1147000 | sporozoite asparagine-rich protein (SLARP) | x | | | 2 | 3 | 1 | 3 | 4 | 6 |
| PF3D7_1216600 | cell traversal protein for ookinetes and sporozoites (CelTOS) | x | | | 0 | 10 | 0 | 4 | 0 | 7 |
| PF3D7_1229100 | Multidrug resistance-associated protein 2 (MRP2) | | | x | 2 | 6 | 1 | 4 | 2 | 7 |
| PF3D7_1243900 | double c2-like domain-containing protein (PfD0C2) | | x | | 8 | 10 | 3 | 6 | 2 | 8 |
| PF3D7_1318300 | Conserved Plasmodium protein, unknown function | | | x | 2 | 4 | 3 | 5 | 0 | 3 |
| PF3D7_1325900 | Conserved Plasmodium protein, unknown function | | | x | 7 | 12 | 2 | 7 | 1 | 9 |
| PF3D7_1335900 | thrombospondin-related anonymous protein (TRAP) | x | | | 0 | 21 | 0 | 20 | 0 | 16 |
| PF3D7_1342500 | sporozoite protein essential for cell traversal (SPECT1) | x | | | 0 | 2 | 0 | 0 | 0 | 1 |
| PF3D7_1349300 | Tyrosine kinase-like protein (TKL3) | | | x | 2 | 10 | 1 | 5 | 1 | 12 |
| PF3D7_1365300 | Conserved Plasmodium protein, unknown function | | | x | 0 | 0 | 1 | 2 | 0 | 4 |
| PF3D7_1405400 | DNA mismatch repair protein, putative | | | x | 0 | 0 | 0 | 0 | 0 | 0 |
| PF3D7_1408700 | Conserved Plasmodium protein, unknown function | | | x | 2 | 7 | 2 | 8 | 4 | 6 |
| PF3D7_1438800 | conservd Plasmodium protein, unkown function | | x | | 0 | 0 | 0 | 0 | 0 | 0 |

| | | | | | | | | | | |
|---|---|---|---|---|---|---|---|---|---|---|
| PF3D7_1465800 | Dynein beta chain, putative | | | x | 3 | 4 | 4 | 6 | 4 | 7 |
| PF3D7_1468100 | conservd Plasmodium protein, unkown function | | x | | 2 | 2 | 2 | 0 | 2 | 4 |
| PF3D7_1469600 | Biotin carboxylase subunit of acetyl CoA carboxylase, putative (ACC) | | | x | 4 | 8 | 0 | 12 | 1 | 12 |

[1]Three sets of genes were chosen for variant identification. The first was a list of 16 genes identified in the literature as potential pre-erythrocytic antigens (Pre-erythrocytic antigens). The other two categories were genes by sera from PfSPZ Vaccine and PfSPZ-CVac vaccinees (PfSPZ Vaccinees, PfSPZ-CVac Vaccinees).

** The large number of SNPs in LSA-1 is possibly a reflection of the difficulty aligning the variable repeat region units in this gene.

**Table S8.** Number of Unique Epitopes When Compared to NF54

| Gene ID | Product Description | Number of Unique Epitopes | | |
|---|---|---|---|---|
| | | 7G8 | NF166.C8 | NF135.C10 |
| PF3D7_0220000 | liver stage antigen 3 (LSA3) | | 3 | 4 |
| PF3D7_0304600 | circumsporozoite protein (CSP) | | 2 | 1 |
| PF3D7_0312400 | pfGSK3 | | | 2 |
| PF3D7_0405300 | liver specific protein 2 (LISP2, sequestrin) | 9 | 20 | 18 |
| PF3D7_0408600 | sporozoite invasion-associated protein 1 (SIAP1) | 5 | 5 | 11 |
| PF3D7_0420000 | zinc finger protein, putative | 9 | 5 | 7 |
| PF3D7_0522400 | conserved Plasmodium protein, unknown function | 1 | 7 | 4 |
| PF3D7_0725100 | conserved Plasmodium membrane protein, unknown function | 6 | | |
| PF3D7_0726400 | conserved Plasmodium membrane protein, unknown function | 8 | 5 | 6 |
| PF3D7_0728600 | zinc finger, C3HC4 type, putative | 2 | | 2 |
| PF3D7_0826100 | E3 ubiquitin-protein ligase, putative | 2 | | 4 |
| PF3D7_0828100 | conserved Plasmodium protein, unkown function | 3 | 3 | 7 |
| PF3D7_0830300 | sporozoite invasion-associated protein-2 (SIAP-2) | 5 | 4 | 5 |
| PF3D7_1021700 | conserved Plasmodium membrane protein, unknown function | 1 | 2 | 1 |
| PF3D7_1030200 | claudin-like apicomplexan microneme protein, putative | 2 | 2 | 2 |
| PF3D7_1035300 | glutamate-rich protein (GLURP) | 2 | | 1 |
| PF3D7_1036400 | liver stage antigen 1 (LSA1) | 3 | 3 | 1 |
| PF3D7_1133400 | apical membrane antigen 1 (AMA1) | 16 | 12 | 15 |
| PF3D7_1138400 | guanylyl cyclase (GCalpha) | 1 | 1 | 1 |

| | | | | |
|---|---|---|---|---|
| PF3D7_1147000 | sporozoite asparagine-rich protein (SLARP) | 1 | 3 | 1 |
| PF3D7_1216600 | cell traversal protein for ookinetes and sporozoites (CelTOS) | | 3 | 1 |
| PF3D7_1229100 | multidrug resistance-associated protein 2 (MRP2) | 9 | 5 | 25 |
| PF3D7_1243900 | double c2-like domain-containing protein (PfD0C2) | 1 | | 3 |
| PF3D7_1318300 | conserved Plasmodium protein, unknown function | 2 | 2 | |
| PF3D7_1335900 | thrombospondin-related anonymous protein (TRAP) | 4 | 4 | 3 |
| PF3D7_1342500 | sporozoite protein essential for cell traversal (SPECT1) | 5 | | 2 |
| PF3D7_1365300 | conserved Plasmodium protein, unknown function | | | 7 |
| PF3D7_1408700 | conserved Plasmodium protein, unknown function | 12 | 14 | 12 |
| PF3D7_1465800 | dynein beta chain, putative | 3 | 7 | 4 |
| PF3D7_1469600 | biotin carboxylase subunit of acetyl CoA carboxylase, putative (ACC) | 5 | 8 | 3 |
| *Number of Unique Epitopes* | | 117 | 121 | 153 |

**Table S9**. Copy number of known drug resistance genes in the four PfSPZ strains

| Gene ID | Product | Drug | NF54 | 7G8 | NF166.C8 | NF135.C10 |
|---|---|---|---|---|---|---|
| PF3D7_0417200 | PfDHFR | sulfadoxine-pyrimethamine | 1 | 1 | 1 | 1 |
| PF3D7_0523000 | PfMDR1 | artesunate, piperaquine, quinine, artemisinin | 1 | 1 | 1 | 4 |
| PF3D7_0709000 | PfCRT | chloroquine | 1 | 1 | 1 | 1 |
| PF3D7_0810800 | PfDHPS | sulfadoxine-pyrimethamine | | | | |
| PF3D7_1224000 | PfGHC1 | sulfadoxine-pyrimethamine | 4 | 2 | 1 | 3 |
| PF3D7_1343700 | PfKELCH13 | artemisinin | | | | |
| PF3D7_1408000 | PfPLASMEPSIN-2 | piperaquine | 1 | 1 | 1 | 1 |
| PF3D7_1408100 | PfPLASMEPSIN-3 | piperaquine | 1 | 1 | 1 | 1 |
| mal_mito_3 | PfCTYB | atovaquone | 1 | 1 | 1 | 1 |

**Table S10.** Non-synonymous SNPs in known drug resistance genes in the four PfSPZ strains (bolded indicates a codon change at that position has previously been shown to be associated with resistance)

| Gene ID | Product | Drug | NF54 | 7G8 | NF166.C8 | NF135.C10 |
|---|---|---|---|---|---|---|
| PF3D7_0417200 | PfDHFR | sulfadoxine-pyrimethamine | - | **N51I**, **S108N** | **N51I**, **C59R**, **S108N** | **N51I**, **C59R**, **S108N**, **I164L** |
| PF3D7_0523000 | PfMDR1 | chloroquine, quinine, mefloquine, amodiaquine, artemisinin | - | **Y184F, S1034C, N1042D, D1246Y** | **Y184F** | F1226Y |
| PF3D7_0709000 | PfCRT | chloroquine | - | **C72S, K76T**, **A220S, N326D, I356L** | - | **M74I, N75E, K76T, A220S, Q271E, N326S, I356T**, **R371I** |
| PF3D7_0810800 | PfDHPS | sulfadoxine-pyrimethamine | - | - | **G437A*** | **S436A**, **K540E** |
| PF3D7_1343700 | PfKELCH13 | artemisinin | - | - | K189T | - |
| mal_mito_3 | PfCYTB | atovaquone | - | - | - | - |

*Reference allele in this case conveys resistance; therefore, NF166.C8 carries the allele conveying resistance, while the other strains match 3D7 in this position (and therefore carry the allele conveying resistance).

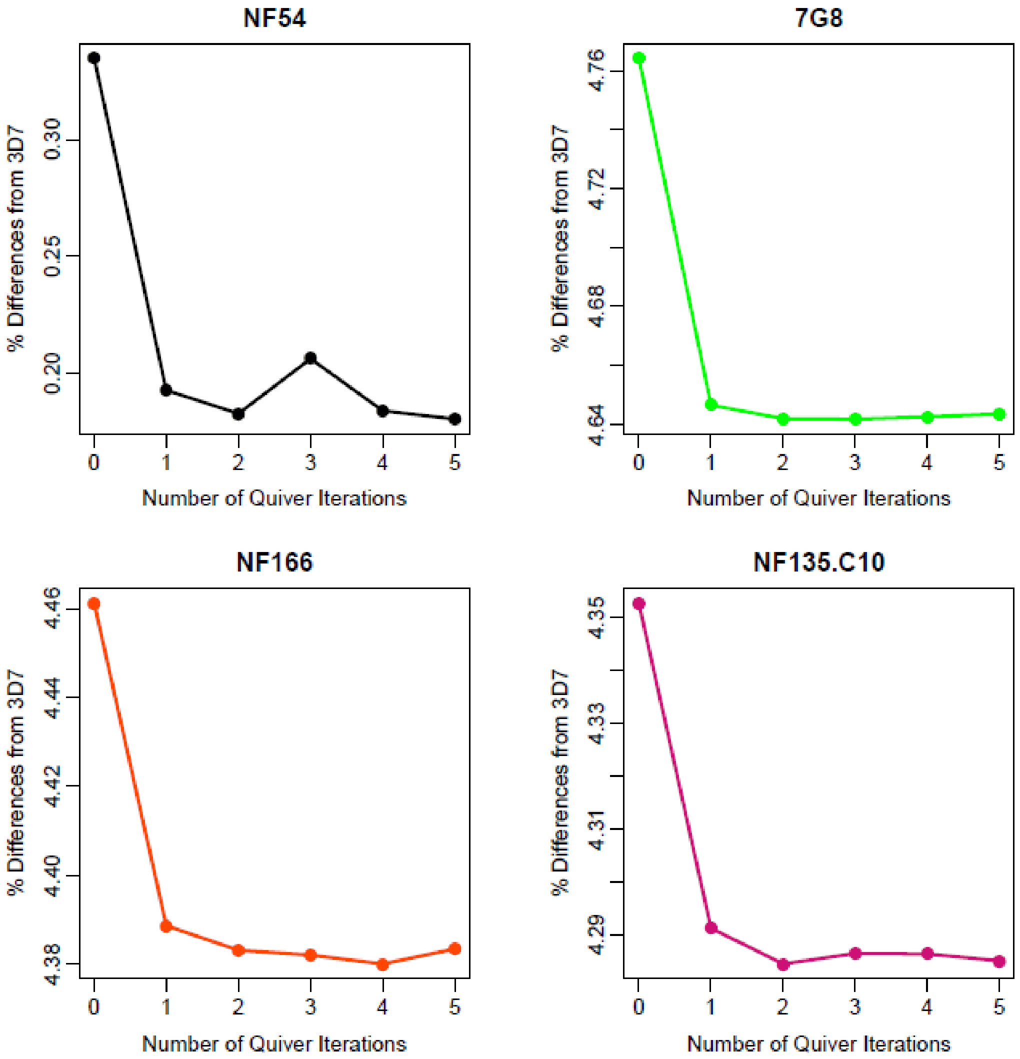


**Figure S1: Polishing PfSPZ assemblies iteratively with Quiver minimized errors in the assemblies.** PacBio and Illumina reads were used to polish the PfSPZ assemblies with Quiver and Pilon, respectively. To minimize the remaining sequencing errors in the assemblies, Quiver was run iteratively on the previously polished version of the assembly, and results are shown for each PfSPZ assembly. The X-axis is the number of Quiver iterations (1-5); the y-axis is the percent difference from the 3D7 reference genome, which includes both base pair differences (such as single-nucleotide polymorphism and indels) and regions of the 3D7 genome that are not present in the PfSPZ assembly.

**Figure S2: *var* exon 1 sequence homologues between 3D7 and the PfSPZ strains**. *var* exon 1 sequences were recovered from each assembly using the ETHA package (see Methods). Shown here are all recovered exon 1 sequences greater than 2.5kb that have an NTS sequence: 45, 66, and 64 for 7G8, NF166.C8, and NF136.C10, respectively. (All exon 1 sequences over 2.5kb in the 7G8 assembly presented here, with or without NTS sequences, are also in the previously published 7G8 assembly from main manuscript reference 33). Each exon 1 sequence in 3D7 is mapped to its identified homolog in NF54 (top left), 7G8 (top right), NF135.C10 (bottom left), and NF166.C8 (bottom right), shown by a ribbon. The color of the ribbon corresponds to percent amino acid identity. Blue: ≤ 0.25, green: ≥ 0.25 and ≤ 0.50, orange: ≥ 0.50 and ≤ 0.75, red: ≥ 0.75. While all 61 exon 1 sequences from 3D7 are present in the NF54 assembly at very high identity, many exon 1 sequences in 7G8, NF166.C8, and NF135.C10 have no identifiable homologs, with the exception of *var2csa* (Pf3D7_1200600).

(See Figure_S2.svg for high resolution image)

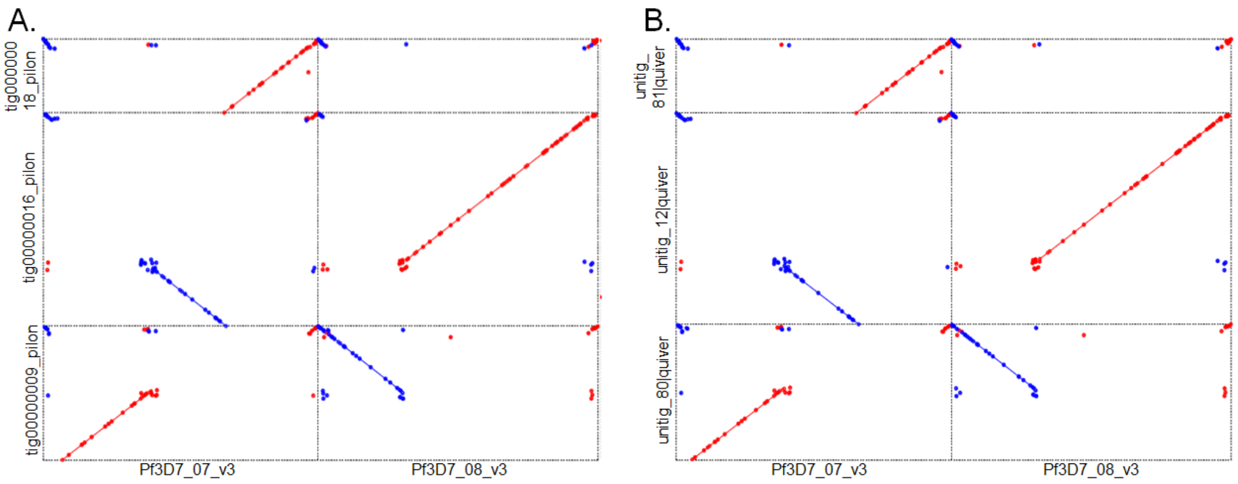


**Figure S3: NF135.C10 contains a chromosomal rearrangement in chromosomes 7 and 8 that is supported by two different assemblers.** Mummerplots of NF135.C10 assembly contigs against the 3D7 reference genome show a translocation and inversion of the end of chromosome 8 and a middle section of chromosome 7. Red alignments indicate an alignment in the same orientation as the 3D7 reference; blue alignments indicate an alignment in the reverse orientation. **A.** Alignments for the Canu assembly, which reconstructs the 3D7 chromosome 7 into two pieces (tig00000009_pilon and tig00000018_pilon) and chromosome 8 in a single piece (tig00000016_pilon), with the end of tig00000016_pilon/Chromosome 8 also aligning in the reverse orientation to the middle section of Chromosome 7 (end of tig00000009_pilon). **B**. An assembly made with the HGAP assembler also shows the same rearrangement. Chromosome 8 is still reconstructed in a single contig (unitg_12|quiver), while chromosome 7 is broken into two contigs (untig_80|quiver and unitig_81|quiver).

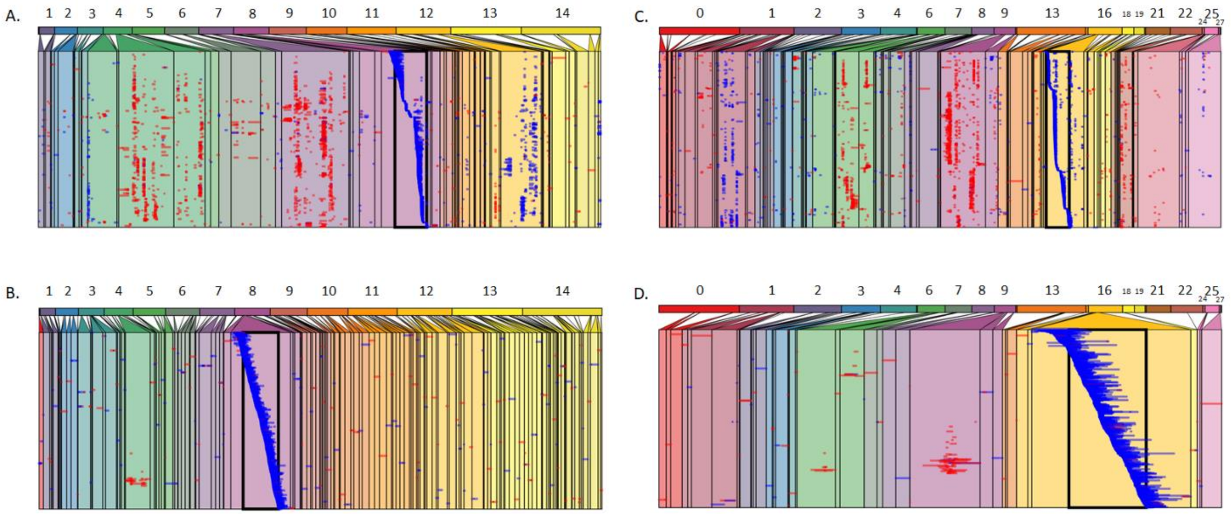


**Figure S4: Validation of chromosomal translocation in NF135.C10 relative to the reference strain 3D7 using cross-mapping of PacBio reads against assemblies.** Assembly results suggest that the genome of NF135.C10 has a chromosomal translocation relative to the reference 3D7 genome. If that is the case, NF135.C10 reads that span the boundary of the translocation region should map partially to the same region in 3D7 (read segment upstream of boundary of translocation) and partly to the region in 3D7 that is the source of the translocated segment (NF135.C10 read segment downstream of the boundary of the translocation). In contrast, our assembly results show that NF166.C8 has a similar genome structure as 3D7 and, therefore, each read from NF166.C8 should map to a single region when aligned against the 3D7 genome.

Panel description: The top track on each of the four panels represents chromosomes in the 3D7 assembly (**A** & **B**) or assembly contigs, which together represent chromosomes in the NF135.C10 assembly (**C** & **D**). The second track shows reads that map to a user-specified region (black box) of a chromosome/contig, which is expanded (zoomed in), with reads ordered according to the 5’ end of the location where they map within this box. The height of the track represents depth of read coverage; regions of those chromosomes/contigs where PacBio reads map in either forward (blue) or reverse (red) orientation are shown. If reads that map to the specified region map only/primarily there, then only that region will be expanded (**B & D**). However, if a segment of each of the same reads maps to a secondary location, that secondary region of the genome is also expanded in the viewer. **A)** NF135.C10 PacBio reads mapped to the 3D7 reference. The black box is 25 Kb wide centered on position 440 Kb of chromosome 8 of 3D7, the proposed boundary of the translocation in the NF135 genome. Segments of these same reads map to other locations, predominantly chromosome 7 (the donor of the translocation in NF135 − see Figure S2), and also regions of chromosomes 4 and 13. **B)** NF166.C8 reads mapped to the same location of chromosome 8 of the 3D7 reference genome. As expected, NF166.C8 reads map to a unique location, showing that the results of panel **A** are not an artifact. **C)** NF166.C8 PacBio reads mapped to contig 16 of the NF135.C10 assembly, the homologous region of 3D7 reference genome shown in **A** and **B**. Similar to panel B, this mapping confirms a different organization of the NF135.C10 genome relative to NF166.C8 and, by proxy, to 3D7. **D)** NF135.C10 PacBio reads mapped to the

NF135.C10 assembly at the region of chromosome 8 homologous to the region of the 3D7 genome shown in **A** and **B**. As expected, these reads map to a single location, showing that the mapping in A is not an artifact of NF135.C10 reads.

A.

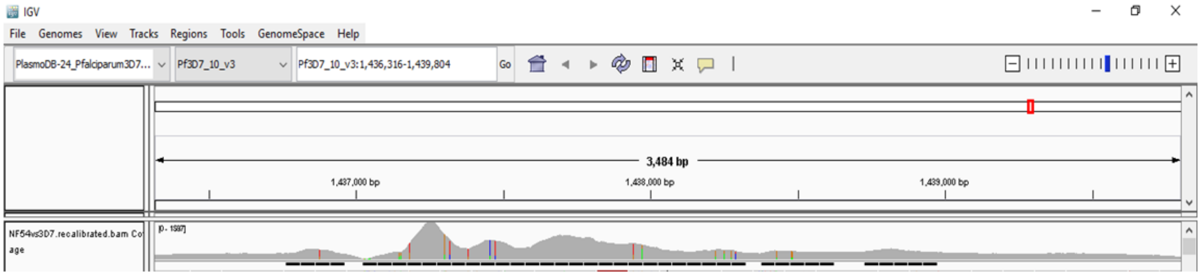


B.

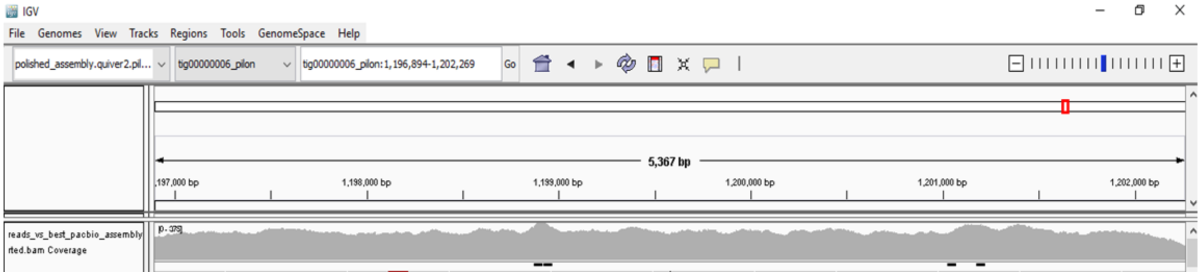


**Figure S5: Illumina read coverage of the repetitive region of liver stage antigen 1 (LSA-1) suggests that the region is longer in NF54 than it is in the 3D7 reference genome. A**. NF54 Illumina reads mapped against the LSA-1 locus on chromosome 10 of the 3D7 reference genome visualized in the Integrative Genome Browser (LSA-1 coordinates: 1,436,316 to 1,439,804 bp). **B.** NF54 Illumina reads mapped against the homologous region in the NF54 assembly (tig00000006_pilon, LSA-1 coordinates: 1,196,860 to 1,202,266 bp).

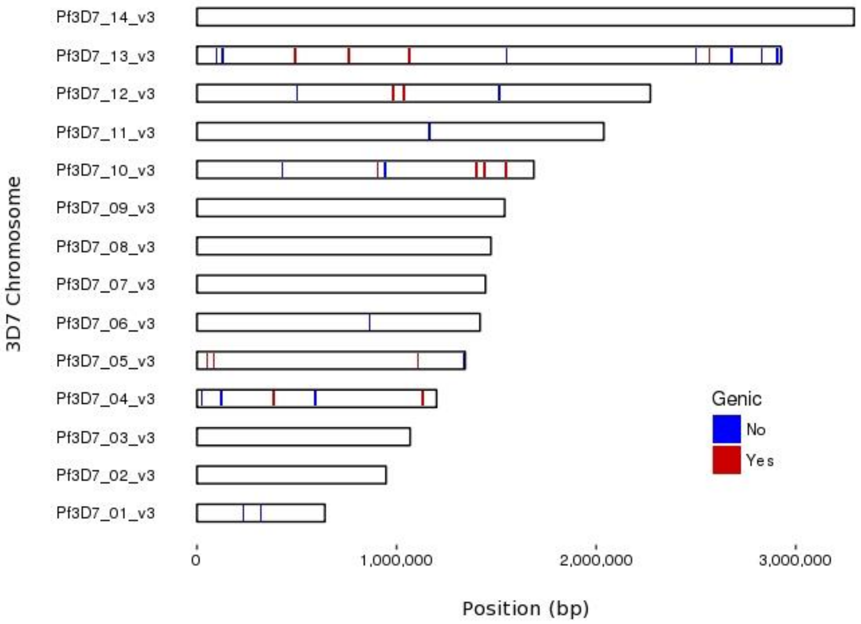


| 3D7 Chromosome | Start Position | Estimated Size | SV Type | Gene Affected |
|---|---|---|---|---|
| Pf3D7_04_v3 | 375,116 | 72 | Deletion | PF3D7_0407600 |
| Pf3D7_04_v3 | 379,839 | 285 | Tandem expansion | PF3D7_0407700 |
| Pf3D7_04_v3 | 1,127,248 | 777 | Tandem expansion | PF3D7_0424900 |
| Pf3D7_05_v3 | 44,357 | 417 | Tandem expansion | PF3D7_0500800 |
| Pf3D7_05_v3 | 78,108 | 126* | Insertion | PF3D7_0501400 |
| Pf3D7_05_v3 | 1,103,711 | 408 | Tandem expansion | PF3D7_0526600 |
| Pf3D7_10_v3 | 899,565 | 189 | Tandem expansion | PF3D7_1021800 |
| Pf3D7_10_v3 | 1,395,621 | 480* | Tandem contraction | PF3D7_1035200 |
| Pf3D7_10_v3 | 1,436,757 | 1,887 | Tandem expansion | PF3D7_1036400 |
| Pf3D7_10_v3 | 1,543,910 | 4,131 | Tandem expansion | PF3D7_1038400 |
| Pf3D7_12_v3 | 974,230 | 1,767 | Tandem expansion | PF3D7_1224000 |
| Pf3D7_12_v3 | 974,433 | 2,043 | Tandem expansion | PF3D7_1224000 |
| Pf3D7_12_v3 | 977,629 | 672 | Tandem expansion | PF3D7_1224100 |
| Pf3D7_12_v3 | 1,030,827 | 327* | Tandem contraction | PF3D7_1225300 |
| Pf3D7_13_v3 | 485,287 | 51 | Tandem expansion | PF3D7_1311300 |
| Pf3D7_13_v3 | 754,733 | 123* | Tandem contraction | PF3D7_1318300 |
| Pf3D7_13_v3 | 1,058,750 | 198 | Tandem expansion | PF3D7_1325400 |
| Pf3D7_13_v3 | 2,561,569 | 681* | Repeat expansion | PF3D7_1363900 |

**Figure S6: Shared structural variants (SVs) affecting base-pair totals between the NF54 and 3D7 PacBio assemblies likely represent errors in the 3D7 reference.** SV errors (indels, deletions, repeat expansions and contractions, and tandem expansions and contractions) were identified by comparing the set of SVs detected in the NF54 assembly and a previously published 3D7 PacBio assembly (main manuscript reference 32). Left: location, by chromosome, of possible errors in the 3D7 reference genome. Thickness of line corresponds to length of the SV error, where red lines indicate SVs affecting genic regions, and blue lines, non-genic regions. (SVs that are very close together may create overlapping vertical lines). Right: List of the potential SV errors that affect genic regions; an asterisk in the estimated size column refers to a SV that had slightly different start or stop coordinates in either the NF54 or 3D7 PacBio assembly, and therefore may represent both errors and true differences between the NF54 and 3D7 genomes (or, alternatively, sequencing or assembly errors on top of an error in the 3D7 reference).

```
PF3D7_1222600    2299 INNDATITFNNPLNNKNVPYQYLNLKDKKQTFQQYNHIKDRENSYPYMHIKGETQQNGMKLKNDINNHHSQNQLSLSKQENNIYKNNVMD
NF54             2299 INNDATITFNNPLNNKNVPYQYLNLKDKKQTFQQYNHIKDRENSYPYMHIKGETQQNGMKLKNDINNHHSQNQLSLSKQENNIYKNNVMD
7G8              2300 INNDATITFNNPLNNKNVPYQYLNLKDKKQTFQQYNHIKDRENSYPYMHIKGETQQNGMKLKNDINNHHSQNQLSLSKQENNIYKNNVMD
NF166            2300 INNDATITFNNPLNNKNVPYQYLNLKDKKQTFQQYNHIKDRENSYPYMHIKGETQQNGMKLKNDINNHHSQNQLSLSKQENNIYKNNVMD
NF135.C10        2303 INNDATITFNNPLNNKNVPYQYLNLKDKKQTFQQYNHIKDRENSYPYMHIKGETQQNGMKLKNDINNHHSQNQLSLSKKENNIYKNNVMD
PfGA01_12002730  2294 INNDATITFNNPLNNKNVPYQYLNLKDKKQTFQQYNHIKDRENSYPYMHIKGETQQNGMKLKNDINNHHSQNQLSLSKQENNIYKNNVMD
PfGN01_12002860  2305 INNDATITFNNPLNNKNVPYQYLNLKDKKQTFQQYNHIKDRENSYPYMHIKGETQQNGMKLKNDINNHHSQNQLSLSKQENNIYKNNVMD
PfCD01_12002750  2304 INNDATITFNNPLNNKNVPYQYLNLKDKKQTFQQYNHIKDRENSYPYMHIKGETQQNGMKLKNDINNHHSQNQLSLSKQENNIYKNNVMD
PfSN01_12002820  2321 INNDATITFNNPLNNKNVPYQYLNLKDKKQTFQQYNHIKDRENSYPYMHIKGETQQNGMKLKNDINNHHSQNQLSLSKQENNIYKNNVMD
PfSD01_12002750  2309 INNDATITFNNPLNNKNVPYQYLNLKDKKQTFQQYNHIKDRENSYPYMHIKGETQQNGMKLKNDINNHHSQNQLSLSKQENNIYKNNVMD
PfML01_12002780  2305 INNDATITFNNPLNNKNVPYQYLNLKDKKQTFQQYNHIKDRENSYPYMHIKGETQQNGMKLKNDINNHHSQNQLSLSKQENNIYKNNVMD
PfGB4_120027200  2304 INNDATITFNNPLNNKNVPYQYLNLKDKKQTFQQYNHIKDRENSYPYMHIKGETQQNGMKLKNDINNHHSQNQLSLSKQENNIYKNNVMD
PfKE01_12002760  2300 INNDATITFNNPLNNKNVPYQYLNLKDKKQTFQQYNHIKDRENSYPYMHIKGETQQNGMKLKNDINNHHSQNQLSLSKQENNIYKNNVMD
PfTG01_12002750  2300 INNDATITFNNPLNNKNVPYQYLNLKDKKQTFQQYNHIKDRENSYPYMHIKGETQQNGMKLKNDINNHHSQNQLSLSKQENNIYKNNVMD
PfHB3_120027400  2290 INNDATITFNNPLNNKNVPYQYLNLKDKKQTFQQYNHIKDRENSYPYMHIKGETQQNGMKLKNDINNHHSQNQLSLSKQENNIYKNNVMD
PfIT_120027700   2299 INNDATITFNNPLNNKNVPYQYLNLKDKKQTFQQYNHIKDRENSYPYMHIKGETQQNGMKLKNDINNHHSQNQLSLSKKENNIYKNNVMD
PfDd2_120027000  2302 INNDATITFNNPLNNKNVPYQYLNLKDKKQTFQQYNHIKDRENSYPYMHIKGETQQNGMKLKNDINNHHSQNQLSLSKKENNIYKNNVMD
PfKH01_12002900  2296 INNDATITFNNPLNNKNVPYQYLNLKDKKQTFQQYNHIKDRENSYPYMHIKGETQQNGMKLKNDINNHHSQNQLSLSKKENNIYKNNVMD
PfKH02_12002760  2295 INNDATITFNNPLNNKNVPYQYLNLKDKKQTFQQYNHIKDRENSYPYMHIKGETQQNGMKLKNDINNHHSQNQLSLSKKENNIYKNNVMD
PPRFG01_1232200  2286 INNDATITFNNPLNNKNVPYQYLNLKDKKQTFQQYNHIKDRENSYPYMHIKGETQQNGMKLKNDINNHHSQNQLSSSNQENNIYKNNVMD
PRCDC_1221900    2231 IDNDATITFNNLLNNKNVPYQYLNVKEKKQTFQQYNHIKDREISYPYMHIKGETQQNSMKLKNDINNHHSQNQLSLSNQENNIYKNNVMD

PF3D7_1222600    2389 EKLLIQNLKACINESNNNN-NNN-------------------------------------------------------------N
NF54             2389 EKLLIQNLKACINESNNNN-NNN-------------------------------------------------------------N
7G8              2390 EKLLIQNLKACINESNNNN-NNN-------------------------------------------------------------N
NF166            2390 EKLLIQNLKACINESNNNNNNNN-------------------------------------------------------------N
NF135.C10        2393 EKLLIQNLKACINESNNNNNNNSSSSSNSNTNGSYKNFITNTNHIRNLDNNSYNANNNFNTQVNSSS*YTPNDNSILNNNNNNNNNNN-N
PfGA01_12002730  2384 EKLLIQNLKACINESNNNN-NNN-------------------------------------------------------------N
PfGN01_12002860  2395 EKLLIQNLKACINESNNNN-NNN-------------------------------------------------------------N
PfCD01_12002750  2394 EKLLIQNLKACINESNNNN-NNN-------------------------------------------------------------N
PfSN01_12002820  2411 EKLLIQNLKACINESNNNN-NNN-------------------------------------------------------------N
PfSD01_12002750  2399 EKLLIQNLKACINESNNNN-NNN-------------------------------------------------------------N
PfML01_12002780  2395 EKLLIQNLKACINESNNNNNNNN-------------------------------------------------------------N
PfGB4_120027200  2394 EKLLIQNLKACINESNNNN-NNN-------------------------------------------------------------N
PfKE01_12002760  2390 EKLLIQNLKACINESNNNN-NNN-------------------------------------------------------------N
PfTG01_12002750  2390 EKLLIQNLKACINESNNNN-NNN-------------------------------------------------------------N
PfHB3_120027400  2380 EKLLIQNLKACINESNNNN-NNN-------------------------------------------------------------N
PfIT_120027700   2389 EKLLIQNLKACINESNNNNNNNSSSSSNSNTNGSYKNFITNTNHIRNLDNNSYNANNNFNTQVNSSS*YTPNDNSILNNNNNNNNNNN-N
PfDd2_120027000  2392 EKLLIQNLKACINESNNNNNNNSSSSSNSNTNGSYKNFITNTNHIRNLDNNSYNANNNFNTQVNSSS*YTPNDNSILNNNNNNNNNNN-N
PfKH01_12002900  2386 EKLLIQNLKACINESNNNNNNNSSSSSNSNTNGSYKNFITNTNHIRNLDNNSYNANNNFNTQVNSSS*YTPNDNSILNNNNNNNNNNN-N
PfKH02_12002760  2385 EKLLIQNLKACINESNNNNNNNSSSSSNSNTNGSYKNFITNTNHIRNLDNNSYNANNNFNTQVNSSS*YTPNDNSILNNNNNNNNNNN-N
PPRFG01_1232200  2376 EKLLIQNLKACINESNNNN-NNSSSSSNSNTNGSYKNFITNTNHIRNLDNNSYNANNNFNTQVNSSSEYTPNDNSILNNNNNNNNNNNNN
PRCDC_1221900    2321 EKLLIQNLQACINESNNNN----SSSSNSNTNGSYKNFLTNTNHIRNLDNNSYNANNNFNTQVNSSSEYTPNDNSILNNNNNNN-----N

PF3D7_1222600    2412 YNVQKRIQNNYYHKGGNNRNI*
NF54             2412 YNVQKRIQNNYYHKGGNNRNI*
7G8              2413 YNVQKRIQNNYYHKGGNNRNI*
NF166            2414 YNVQKRIQNNYYHKGGNNRNI*
NF135.C10        2481 YNVQKRIQNNYYHKGGNNRNI*
PfGA01_12002730  2407 YNVQKRIQNNYYHKGGNNRNI*
PfGN01_12002860  2418 YNVQKRIQNNYYHKGGNNRNI*
PfCD01_12002750  2417 YNVQKRIQNNYYHKGGNNRNI*
PfSN01_12002820  2434 YNVQKRIQNNYYHKGGNNRNI*
PfSD01_12002750  2422 YNVQKRIQNNYYHKGGNNRNI*
PfML01_12002780  2419 YNVQKRIQNNYYHKGGNNRNI*
PfGB4_120027200  2417 YNVQKRIQNNYYHKGGNNRNI*
PfKE01_12002760  2413 YNVQKRIQNNYYHKGGNNRNI*
PfTG01_12002750  2413 YNVQKRIQNNYYHKGGNNRNI*
PfHB3_120027400  2403 YNVQKRIQNNYYHKGGNNRNI*
PfIT_120027700   2477 YNVQKRIQNNYYHKGGNNRNI*
PfDd2_120027000  2480 YNVQKRIQNNYYHKGGNNRNI*
PfKH01_12002900  2474 YNVQKRIQNNYYHKGGNNRNI*
PfKH02_12002760  2473 YNVQKRIQNNYYHKGGNNRNI*
PPRFG01_1232200  2465 YNVQKRIQNNYYHKGGNNRNI*
PRCDC_1221900    2402 YNMQKRIQNNYYHKGGNNRNI*
```

**Figure S7: An indel in the 3' end of AP2-G (PF1222600) appears to be specific to isolates from Southeast Asia (including the heterologous CHMI clone NF135.C10).** The alignment above shows the 3'end of the AP2-G gene. 3D7 and the four PfSPZ clones are shown, followed by the AP2-G sequences from previously published long-read assemblies (reference 33 in main manuscript) and two non-human primate *Plasmodium* species (PPRFG01: *P. praefalciparum*; PRCDC: *P. reichenowi*). Sequences from the latter two groups were pulled from PlasmoDBv41. A premature stop codon (yellow highlighting) in a ~68 amino acid insertion (relative to 3D7) in isolates of known or suspected Southeast Asian origin (bold text) is shown. PfIT, while originally sampled from South America, has previously been shown to have signs of contamination, sharing genetic singles with isolates from Southeast Asia (supplemental reference 1).

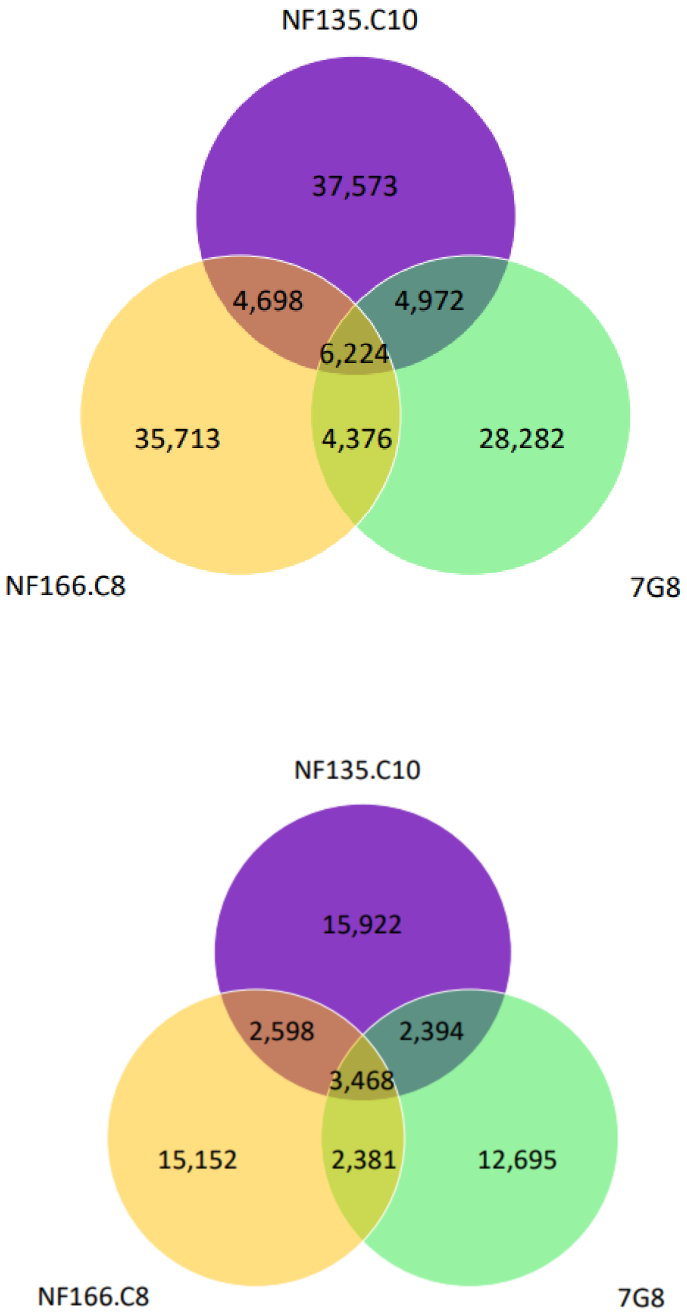


**Figure 8: Shared and unique SNPs between the four PfSPZ strains.** Genome-wide (top) and restricted to non-synonymous SNPs (bottom). SNPs were detected by aligning each genome to the 3D7 genome with nucmer and nucmer's show-snps to detect variants. Unique and shared SNPs were then characterized by comparing the positions of SNPs in each heterologous CHMI dataset.

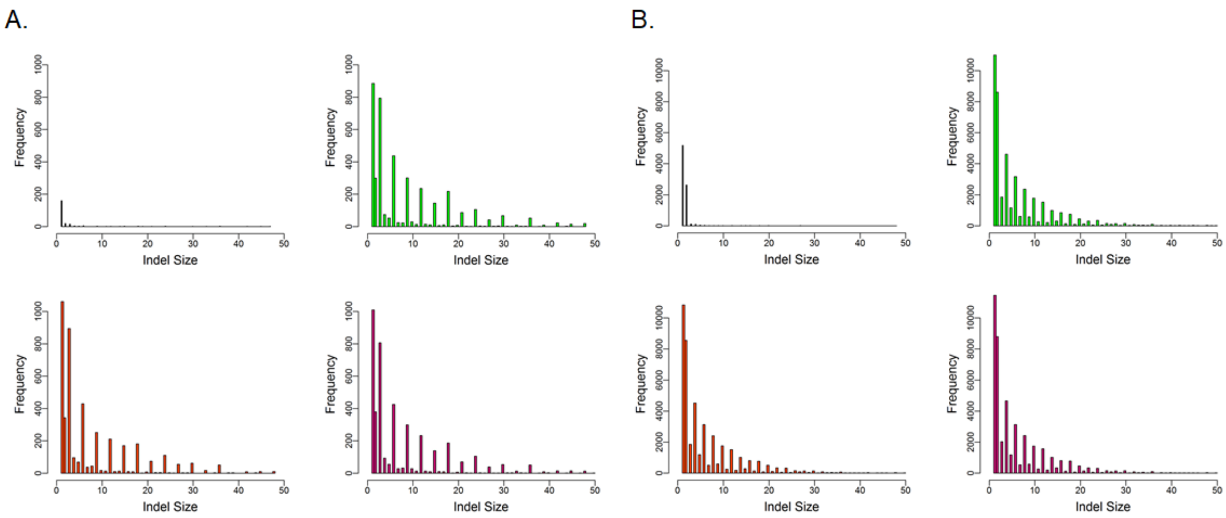


**Figure S9: Distribution of indels.** Small indels (insertions and deletions < 50 bp in size) were identified in each assembly. **A.** Histograms of indels in each assembly in coding regions show that indels tend to be a size in length of multiples-of-three, although some non-multiples-of-three occur (particularly single-bp indels) possibly representing remaining sequencing error). **B.** Histograms of indels from non-coding regions in each assembly show that multiples-of-two indels are more common. NF54 (black), 7G8 (green), NF166.C8 (orange), and NF135.C10 (hot pink).

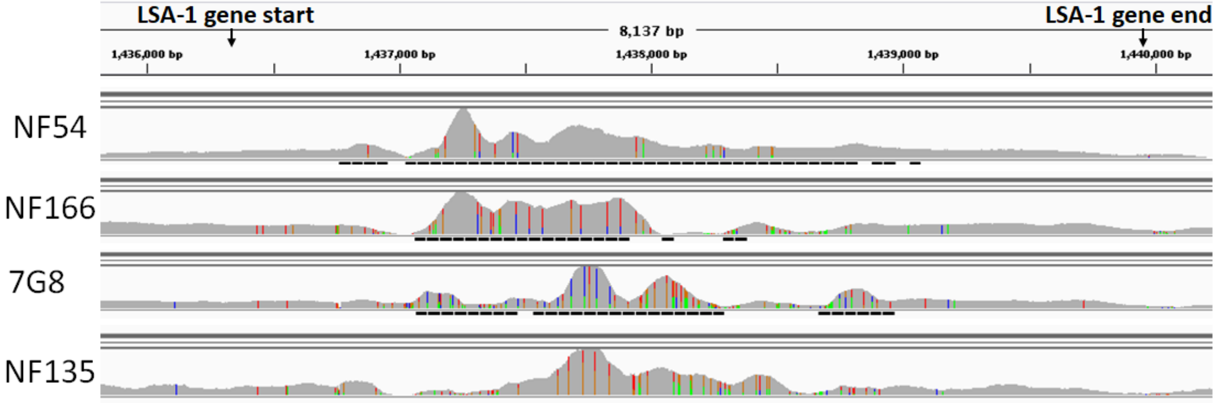


**Figure S10: The repeat region of LSA-1 is variable in the four PfSPZ strains.** Mapping Illumina reads from each PfSPZ strain against the 3D7 reference revealed a pile-up of reads on chromosome 10 in a region containing liver stage antigen 1 (LSA-1) (visualized here in Integrative Genome Browser; LSA-1 coordinates: 1,436,316 to 1,439,804 bp) as was shown between NF54 and 3D7. The differences in read depth and coverage of this region also reflect variations in the size of the repeat region of LSA-1

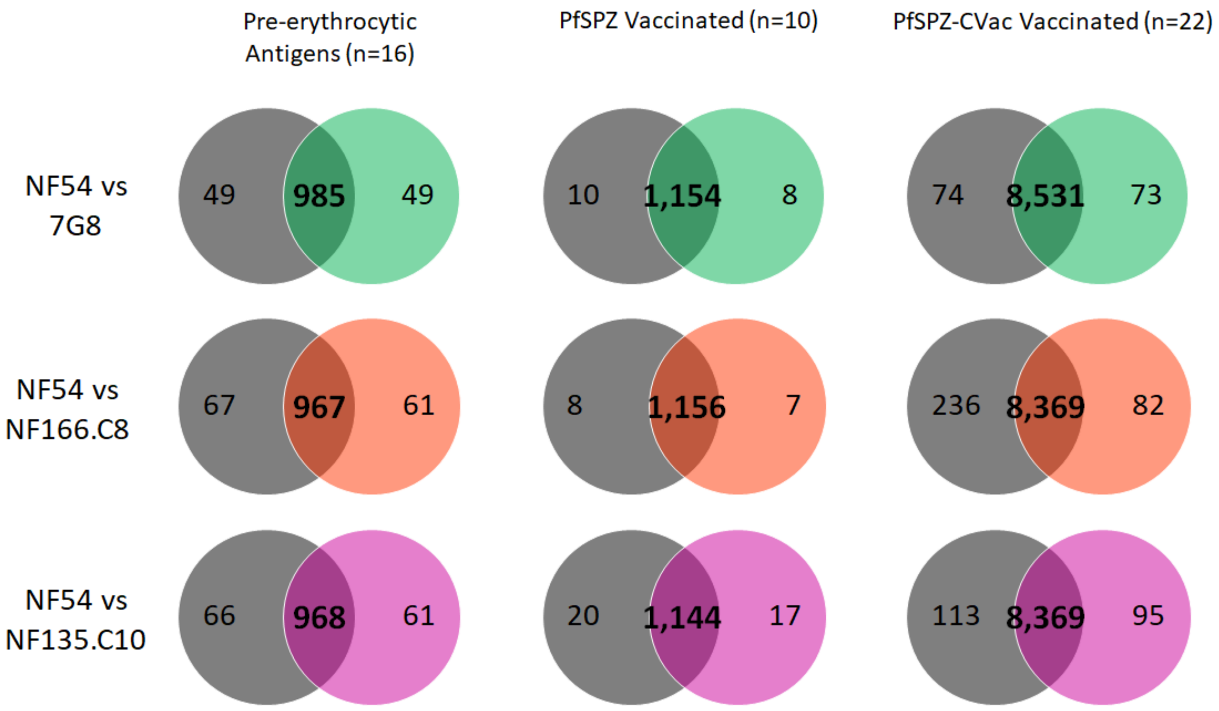


**Figure S11: Shared MHC class 1 epitope sequences between NF54 and the three heterologous CHMI strains in 42 genes of interest.** Pre-erythrocytic genes of interest were identified either from a review of the literature (first column), or genes whose products were recognized by sera from vaccinees of whole organism malaria vaccine trials in PfSPZ (second column) and PfSPZ-CVac (third column). (While these genes were recognized with B cell approaches, many important pre-erythrocytic antigens, such as CSP, have been shown to have both B and T cell epitopes present). For a full list of genes that fall into each of the three categories, refer to Table S6. Shared epitopes between NF54 (black) and 7G8 (green), NF166.C8 (orange), and NF135.C10 (hot pink) are shown in the center of the Venn diagrams, while epitope sequences unique to NF54 or the other heterologous PfSPZ strain are shown in the circles to the left and right.

**Figure S12: Predicted CD8+ T cell epitopes, by position, in selected pre-erythrocytic genes of interest.**

See attached pdf (Figure_S12.pdf)

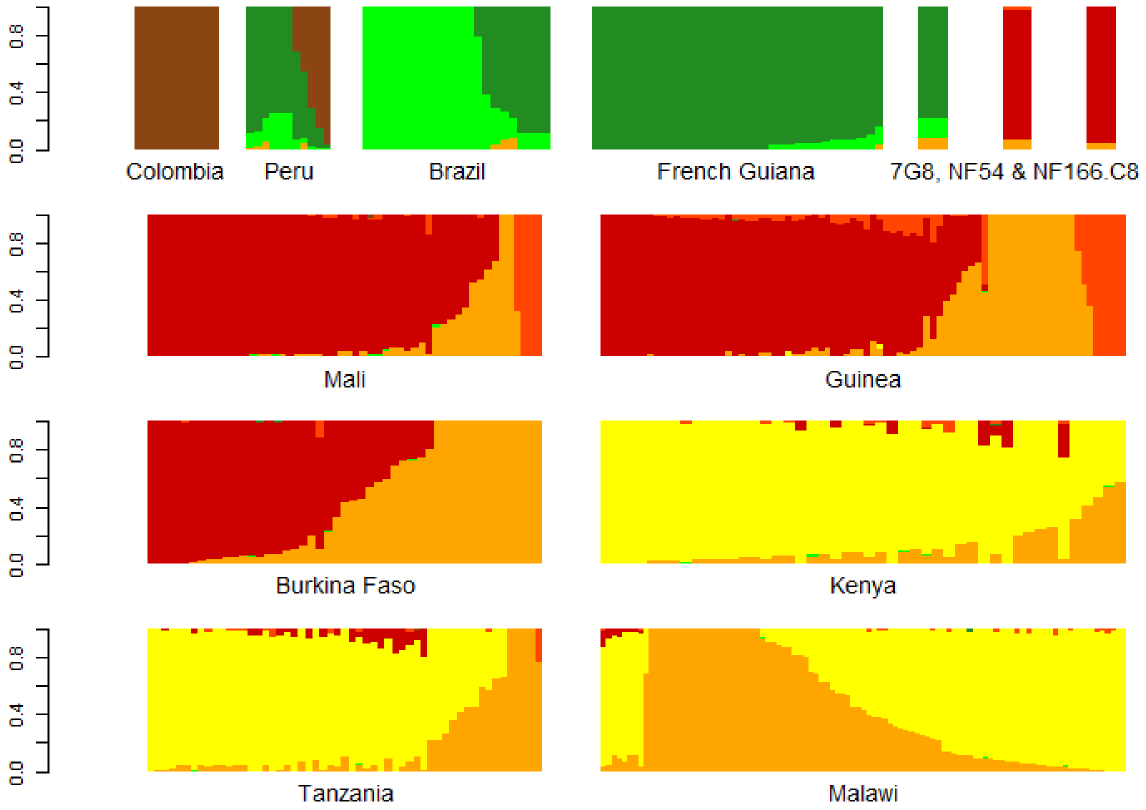


**Figure S13:** Admixture analysis of South American and African clinical isolates place NF54, 7G8, and NF166.C8 amongst clinical isolates from their respective geographic origins. Admixture analysis was done using 16,802 biallelic variable positions identified in 461 isolates from South America and Africa, along with NF54, NF166.C8, and 7G8, identifying *K*=7 subpopulations. Each color represents one of seven subpopulations; each column of a plot is a sample, with the height of each bar representing the proportion of the genome that was assigned by the model to one of the *K* subpopulations. (Single bars representing 7G8, NF54, and NF166.C8 have been enlarged to aid visualization.)
