## Supplemental Figure 2 for "Strains used in whole organism *Plasmodium falciparum* vaccine trials differ in genome structure, sequence, and immunogenic potential"

PF3D7.0100300.1

PF3D7.0200100.1

PF3D7.0223500.1

PF3D7.0300100.1

PF3D7.0400100.1

PF3D7.0412700.1

PF3D7.0412900.1

PF3D7.0420700.1

PF3D7.0420900.1

PF3D7.0421300.1

PF3D7.0426000.1

PF3D7.0500100.1

PF3D7.0600200.1

PF3D7.0600400.1

PF3D7.0617400.1

PF3D7.0700100.1

PF3D7.0711700.1

PF3D7.0712000.1

PF3D7.0712400.1

PF3D7.0712600.1

PF3D7.0733000.1

PF3D7.0800100.1

PF3D7.0800200.1

PF3D7.0800300.1

PF3D7.0808700.1

PF3D7.0809100.1

PF3D7.0833500.1

PF3D7.0937600.1

PF3D7.1000100.1

PF3D7.1041300.1

PF3D7.1100100.1

PF3D7.1150400.1

PF3D7.1200100.1

PF3D7.1219300.1

PF3D7.1240400.1

PF3D7.1240600.1

PF3D7.1255200.1

PF3D7.1300100.1

var7G8.08n04

var7G8.01n03

var7G8.n.add

var7G8.07n01

var7G8.05n04

PF3D7.1373500.1

var7G8.01n02

PF3D7.0100100.1

var7G8.04n04

PF3D7.1300300.1

var7G8.09n02

PF3D7.1240300.1

var7G8.05n05

var7G8.07n05

var7G8.12n09

var7G8.04n02

var7G8.05n01

var7G8.03n01

var7G8.12n02

var7G8.08n05

PF3D7.0412400.1

var7G8.04n03

var7G8.08n02

var7G8.07n06

var7G8.06n01

var7G8.12n04

PF3D7.0425800.1

var7G8.05n02

var7G8.05n06

PF3D7.0400400.1

var7G8.01n01

var7G8.14n01

PF3D7.0115700.1

var7G8.08n01

var7G8.12n06

PF3D7.0900100.1

var7G8.12n03

var7G8.09n01

PF3D7.1240900.1

var7G8.02n01

var7G8.12n07

var7G8.01n04

var7G8.08n03

PF3D7.0712800.1

PF3D7.0712900.1

var7G8.07n07

var7G8.07n02

PF3D7.0712300.1

var7G8.07n04

PF3D7.0937800.1

var7G8.07n03

var7G8.04n05

PF3D7.0808600.1

PF3D7.0632800.1

var7G8.12n05

var7G8.12n01

PF3D7.1100200.1

var7G8.05n03

PF3D7.1200600.1

var7G8.04n01

PF3D7.0324900.1

var7G8.12n10

PF3D7.0632500.1

var7G8.12n08

PF3D7.1200400.1

PF3D7.0421100.1

PF3D7.0413100.1

var7G8.12n12

PF3D7.0100300.1

PF3D7.0200100.1

PF3D7.0300100.1

PF3D7.0400100.1

PF3D7.0412900.1

PF3D7.0413100.1

PF3D7.0421100.1

PF3D7.0421300.1

PF3D7.0426000.1

PF3D7.0600400.1

PF3D7.0617400.1

PF3D7.0711700.1

PF3D7.0712300.1

PF3D7.0712400.1

PF3D7.0712800.1

PF3D7.0733000.1

PF3D7.0800100.1

PF3D7.0800300.1

PF3D7.0808700.1

PF3D7.0833500.1

PF3D7.1100200.1

PF3D7.1219300.1

PF3D7.1240400.1

PF3D7.1240600.1

PF3D7.1300100.1

PF3D7.1373500.1

PF3D7.0632500.1

NF135.scn59

NF135.uni0036.1

NF135.uni0039.1

NF135.uni0082.1

NF135.scn51

NF135.scn47

PF3D7.0712000.1

PF3D7.0712900.1

NF135.scn36

NF135.scn23

NF135.scn52

NF135.scn26

PF3D7.0324900.1

NF135.scn49

NF135.scn29

NF135.scn15

PF3D7.0400400.1

PF3D7.1200100.1

NF135.scn14

NF135.uni0064.1

NF135.scn44

NF135.uni0067.1

NF135.uni0066.1

PF3D7.0115700.1

NF135.uni0054.1

NF135.scn25

PF3D7.1240900.1

PF3D7.0425800.1

NF135.scn43

NF135.uni0068.1

NF135.uni0081.1

NF135.scn35

PF3D7.0412700.1

PF3D7.0420700.1

NF135.uni0053.1

PF3D7.0420900.1

NF135.uni0055.1

NF135.scn37

NF135.scn41

NF135.scn61

PF3D7.1300300.1

NF135.uni0063.1

PF3D7.0712600.1

PF3D7.0700100.1

NF135.scn28

NF135.uni0020.1

NF135.uni0052.1

PF3D7.0100100.1

NF135.scn57

NF135.scn45

NF135.uni0056.1

NF135.scn32

NF135.uni0078.1

PF3D7.0808600.1

NF135.uni0061.1

PF3D7.1255200.1

NF135.uni0073.1

PF3D7.1200400.1

NF135.scn10

NF135.uni0051.1

NF135.scn34

PF3D7.1041300.1

NF135.uni0029.1

PF3D7.0600200.1

NF135.scn3

NF135.uni0074.1

PF3D7.0500100.1

NF135.scn31

PF3D7.0809100.1

NF135.scn19

PF3D7.0632800.1

PF3D7.1200600.1

NF135.uni0027.1

NF135.uni0062.1

NF135.uni0079.1

PF3D7.1240300.1

NF135.uni0071.1

PF3D7.1000100.1

PF3D7.0800200.1

NF135.uni0077.1

PF3D7.0937800.1

NF135.scn27

PF3D7.1150400.1

PF3D7.0937600.1

NF135.scn11

NF135.scn33

PF3D7.0223500.1

NF135.scn13

NF135.uni0065.1

PF3D7.1100100.1

PF3D7.0900100.1

NF135.uni0060.1

PF3D7.0412400.1

NF135.uni0076.1

NF135.uni0075.1

NF135.uni0058.1

NF135.scn58

NF135.scn8

PF3D7.0100300.1

PF3D7.0200100.1

PF3D7.0300100.1

PF3D7.0400100.1

PF3D7.0412900.1

PF3D7.0413100.1

PF3D7.0421100.1

PF3D7.0426000.1

PF3D7.0700100.1

PF3D7.0711700.1

PF3D7.0712000.1

PF3D7.0712600.1

PF3D7.0800200.1

PF3D7.0800300.1

PF3D7.0808700.1

PF3D7.0809100.1

PF3D7.0833500.1

PF3D7.0900100.1

PF3D7.0937800.1

PF3D7.1000100.1

PF3D7.1200100.1

PF3D7.1219300.1

PF3D7.1240300.1

PF3D7.1240600.1

PF3D7.1255200.1

PF3D7.1300100.1

NF166.scn32

PF3D7.0412400.1

NF166.uni0057.1

NF166.scn9

NF166.uni0065.1

NF166.uni0060.1

NF166.uni0072.1

NF166.uni0062.1

PF3D7.0600200.1

PF3D7.1373500.1

NF166.scn60

PF3D7.0600400.1

NF166.scn55

PF3D7.0500100.1

PF3D7.0425800.1

NF166.uni0044.1

NF166.scn10

NF166.uni0067.1

PF3D7.0632500.1

NF166.scn54

NF166.scn53

NF166.uni0071.1

NF166.scn11

NF166.uni0064.1

NF166.scn65

NF166.uni0061.1

PF3D7.1200400.1

NF166.uni0023.1

NF166.scn15

NF166.uni0050.1

NF166.scn22

PF3D7.1100100.1

NF166.uni0053.1

NF166.scn38

NF166.scn34

NF166.uni0049.1

NF166.scn69

PF3D7.0421300.1

NF166.uni0054.1

PF3D7.0412700.1

NF166.uni0052.1

NF166.uni0051.1

PF3D7.0115700.1

NF166.scn12

PF3D7.0808600.1

NF166.scn40

NF166.scn1

NF166.scn46

PF3D7.0632800.1

NF166.scn52

NF166.scn62

PF3D7.0712300.1

NF166.uni0056.1

NF166.scn45

PF3D7.0712400.1

NF166.uni0007.1

NF166.uni0055.1

PF3D7.0712900.1

NF166.scn24

PF3D7.0733000.1

NF166.scn44

NF166.uni0041.1

NF166.uni0042.1

NF166.uni0046.1

PF3D7.1041300.1

PF3D7.0937600.1

NF166.scn35

NF166.uni0073.1

NF166.uni0069.1

PF3D7.1150400.1

NF166.uni0070.1

NF166.uni0043.1

PF3D7.0100100.1

NF166.scn5

NF166.scn59

PF3D7.1240900.1

PF3D7.1200600.1

NF166.uni0048.1

PF3D7.0324900.1

NF166.uni0026.1

PF3D7.0617400.1

NF166.uni0059.1

NF166.uni0068.1

PF3D7.0800100.1

NF166.uni0047.1

PF3D7.0400400.1

PF3D7.0420700.1

NF166.scn63

NF166.scn20

PF3D7.0223500.1

NF166.scn39

PF3D7.0420900.1

PF3D7.1240400.1

NF166.scn42

PF3D7.0712800.1

NF166.scn58

NF166.scn43

PF3D7.1300300.1

PF3D7.1100200.1

NF166.scn17

NF166.scn26

NF54.uni0076.1

NF54.uni0017.1

NF54.uni0026.1

PF3D7.0425800.1

NF54.uni0070.1

NF54.uni0016.1

PF3D7.0712800.1

NF54.uni0005.1

NF54.uni0024.1

PF3D7.0800100.1

NF54.uni0039.1

NF54.uni0038.1

PF3D7.1240600.1

NF54.uni0029.1

NF54.uni0015.1

PF3D7.1373500.1

NF54.uni0027.1

tig00000014.pilon.1

PF3D7.0412900.1

PF3D7.0421100.1

NF54.uni0004.1

PF3D7.0712900.1

NF54.uni0014.1

PF3D7.1200400.1

NF54.uni0019.1

PF3D7.0420900.1

NF54.uni0020.1

NF54.uni0018.1

NF54.uni0021.1

PF3D7.0937600.1

PF3D7.0100300.1

NF54.uni0022.1

NF54.uni0023.1

PF3D7.0600400.1

NF54.uni0052.1

PF3D7.0426000.1

NF54.uni0028.1

PF3D7.0712600.1

NF54.uni0025.1

PF3D7.0200100.1

NF54.uni0031.1

PF3D7.0711700.1

PF3D7.1100100.1

PF3D7.0808700.1

NF54.uni0033.1

NF54.uni0035.1

NF54.uni0034.1

PF3D7.0712300.1

PF3D7.0412700.1

PF3D7.0223500.1

NF54.uni0037.1

PF3D7.0712400.1

PF3D7.1200100.1

NF54.uni0040.1

NF54.uni0041.1

PF3D7.0809100.1

NF54.uni0049.1

PF3D7.0420700.1

NF54.uni0043.1

PF3D7.1300100.1

NF54.uni0044.1

PF3D7.0324900.1

NF54.uni0045.1

PF3D7.0100100.1

NF54.uni0046.1

PF3D7.1300300.1

NF54.uni0047.1

PF3D7.0632800.1

NF54.uni0050.1

PF3D7.0412400.1

NF54.uni0048.1

PF3D7.0115700.1

NF54.uni0042.1

PF3D7.0700100.1

NF54.uni0032.1

NF54.uni0051.1

NF54.uni0030.1

PF3D7.0300100.1

NF54.uni0053.1

PF3D7.0808600.1

NF54.uni0054.1

PF3D7.0500100.1

NF54.uni0055.1

PF3D7.1041300.1

NF54.uni0056.1

PF3D7.0937800.1

NF54.uni0057.1

PF3D7.0413100.1

NF54.uni0060.1

PF3D7.0833500.1

NF54.uni0059.1

PF3D7.1219300.1

NF54.uni0061.1

PF3D7.1240400.1

NF54.uni0058.1

PF3D7.0900100.1

NF54.uni0065.1

PF3D7.1240900.1

NF54.uni0063.1

PF3D7.1255200.1

NF54.uni0064.1

PF3D7.0712000.1

NF54.uni0062.1

PF3D7.1000100.1

NF54.uni0066.1

PF3D7.0400100.1

PF3D7.0733000.1

NF54.uni0067.1

NF54.uni0068.1

PF3D7.1240300.1

NF54.uni0069.1

PF3D7.0617400.1

NF54.uni0072.1

PF3D7.0600200.1

PF3D7.0421300.1

PF3D7.1200600.1

NF54.uni0073.1

PF3D7.0800300.1

NF54.uni0074.1

PF3D7.1100200.1

NF54.uni0075.1

PF3D7.0800200.1

NF54.uni0071.1

PF3D7.1150400.1

NF54.uni0077.1

PF3D7.0400400.1

NF54.uni0078.1

PF3D7.0632500.1
