## Supplemental Figure 12 for "Strains used in whole organism *Plasmodium falciparum* vaccine trials differ in genome structure, sequence, and immunogenic potential"

### PF3D7_0206900_2 / MSP5 (261 aa)

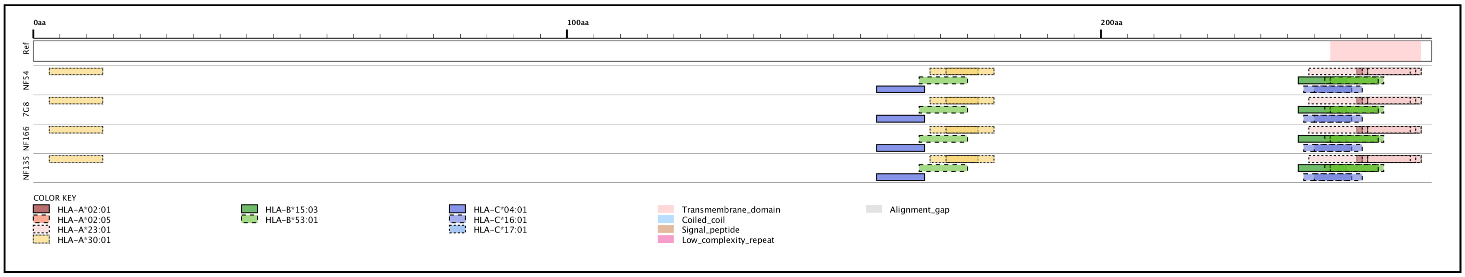


### PF3D7_0207000 / MSP4 (272 aa)

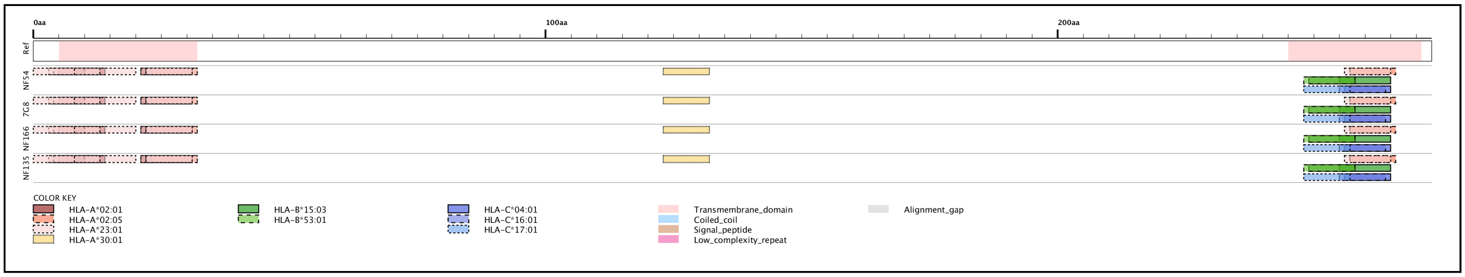


### PF3D7_0220000 / LSA3 (1559 aa)

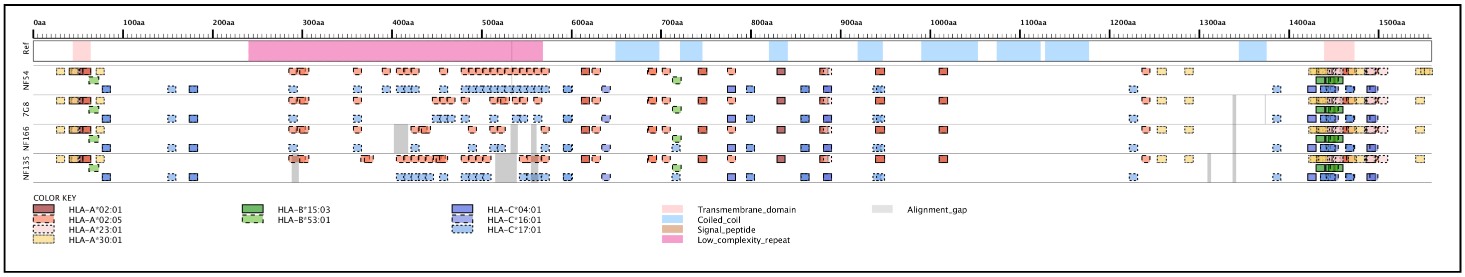


### PF3D7_0304600 / CSP (401 aa)

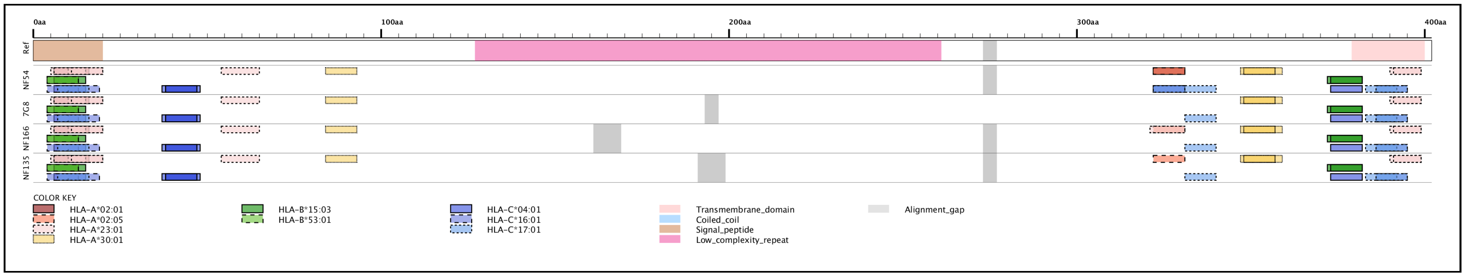


### PF3D7_0405300 / LISP2 (1999 aa)

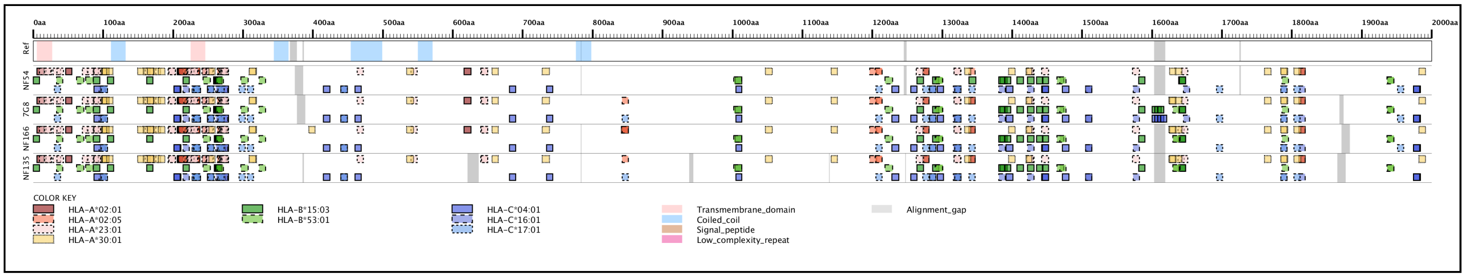


### PF3D7_0408600 / SIAP1 (984 aa)

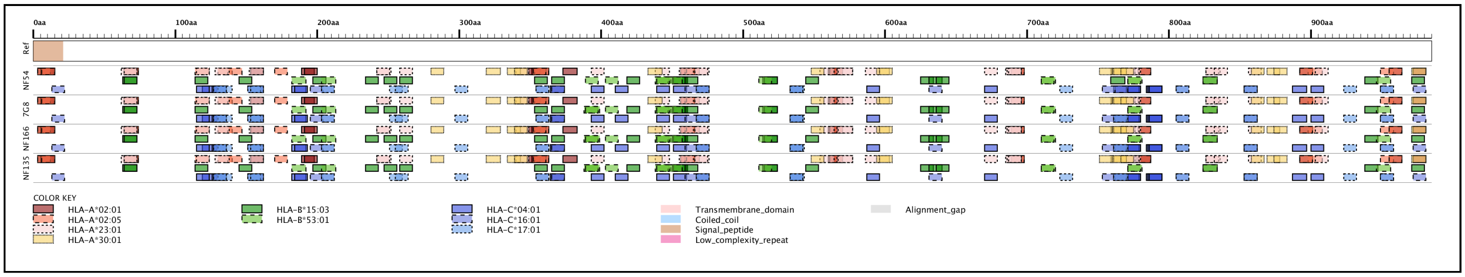


### PF3D7_0408700 / SPLP1/SPECT2 (845 aa)

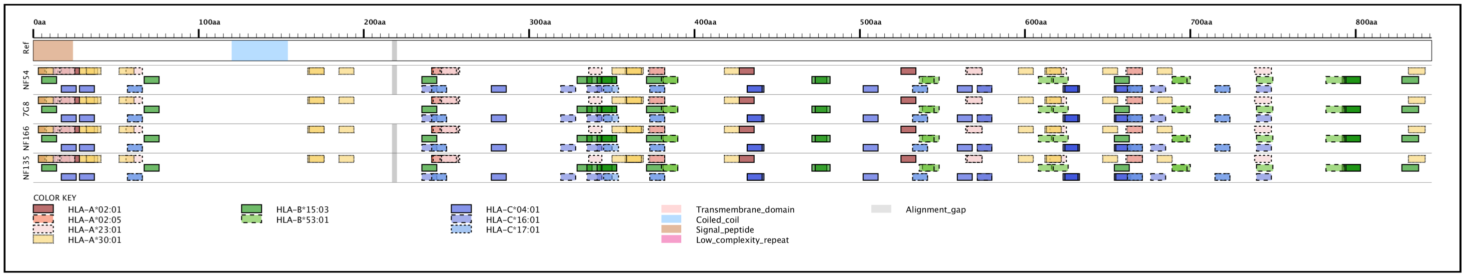


### PF3D7_0812300 / SSP3 (461 aa)

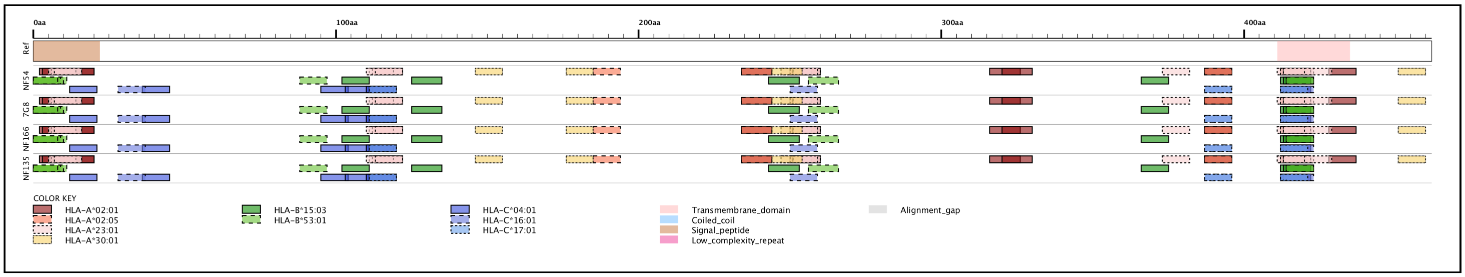


### PF3D7_0830300 / SIAP2 (388 aa)

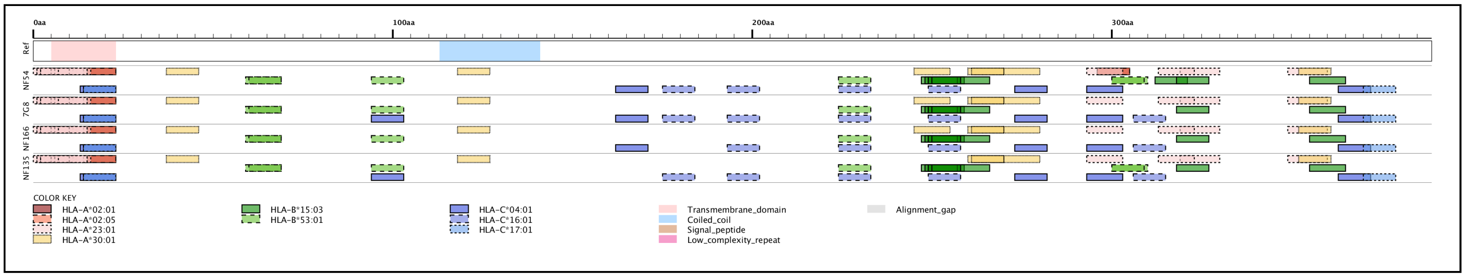


### PF3D7_1036400 / LSA1 (1802 aa)

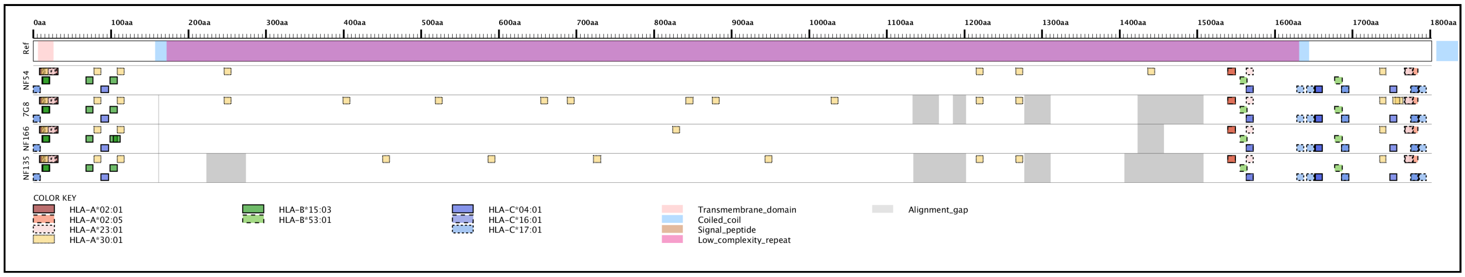


### PF3D7_1121600 / EXP1 (162 aa)

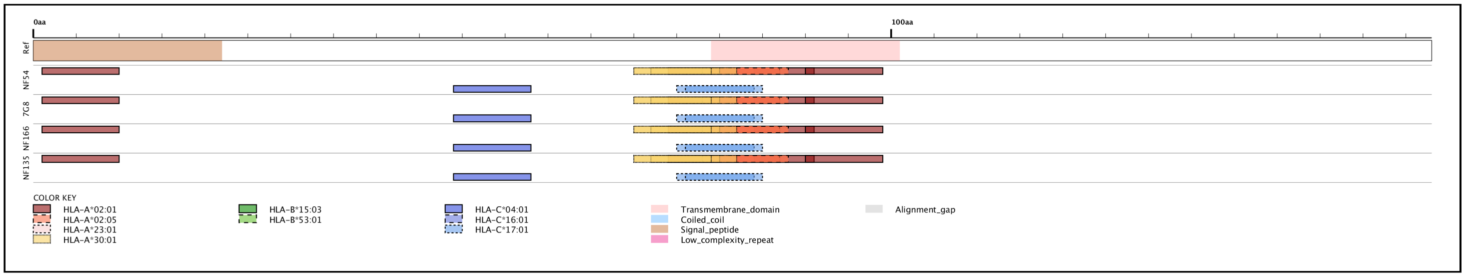


### PF3D7_1133400 / AMA1 (622 aa)

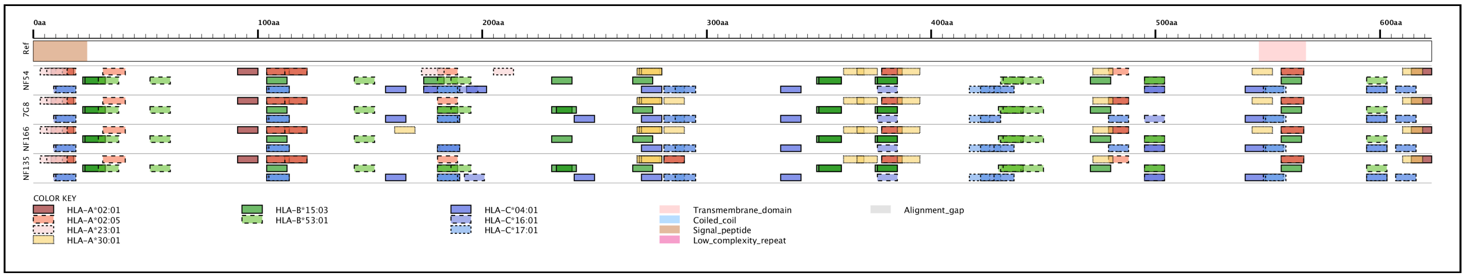


### PF3D7_1147000 / SLARP (2963 aa)

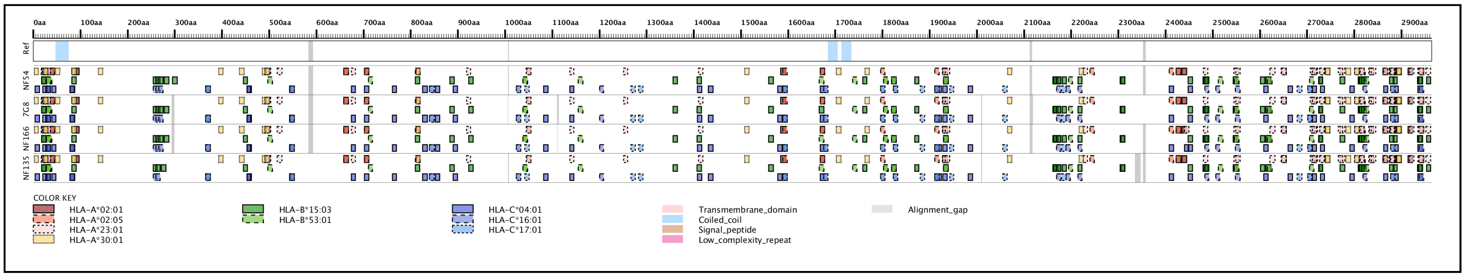


### PF3D7_1216600 / CelTOS (182 aa)

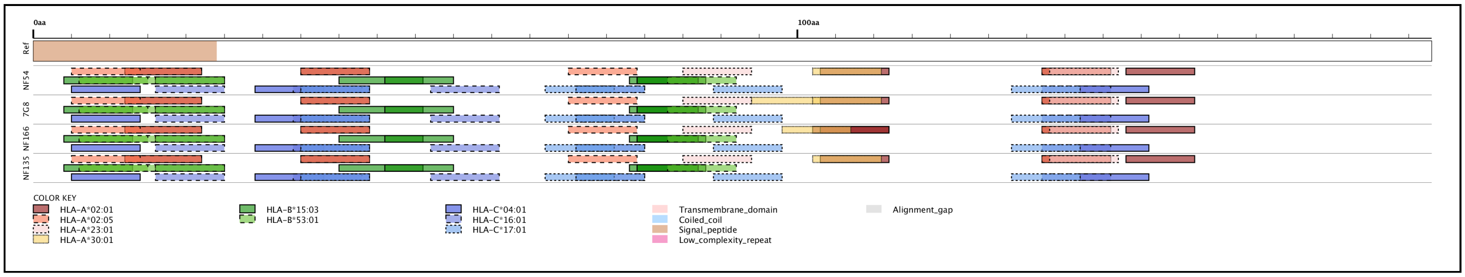


### PF3D7_1335900 / TRAP (574 aa)

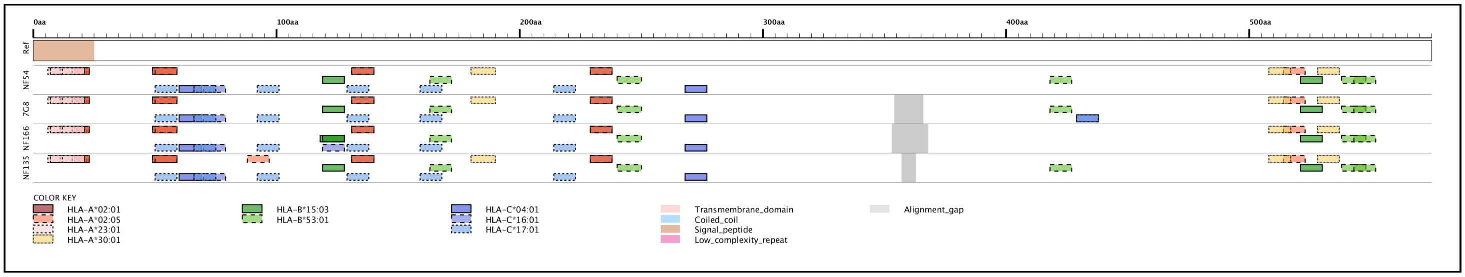


### PF3D7_1342500 / SPECT1 (245 aa)

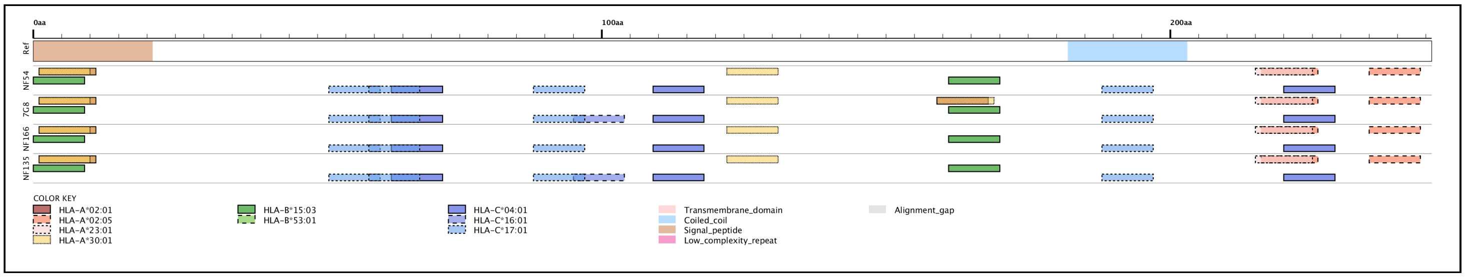
